# Interferon induced circRNAs escape herpesvirus host shutoff and suppress lytic infection

**DOI:** 10.1101/2023.09.07.556698

**Authors:** Sarah E. Dremel, Takanobu Tagawa, Vishal N. Koparde, Jesse H. Arbuckle, Thomas M. Kristie, Laurie T. Krug, Joseph M. Ziegelbauer

## Abstract

A first line of defense during infection is expression of interferon (IFN)-stimulated gene products which suppress viral lytic infection. To combat this, herpesviruses express endoribonucleases to deplete host RNAs. Here we demonstrate that IFN-induced circular RNAs (circRNAs) can escape viral-mediated degradation. We performed comparative circRNA expression profiling for representative alpha- (Herpes simplex virus-1, HSV-1), beta- (human cytomegalovirus, HCMV), and gamma-herpesviruses (Kaposi sarcoma herpesvirus, KSHV; murine gamma-herpesvirus 68, MHV68). Strikingly, we found that circRNAs are, as a population, resistant to host shutoff. This observation was confirmed by ectopic expression assays of human and murine herpesvirus endoribonucleases. During primary lytic infection, ten circRNAs were commonly regulated across all subfamilies of human herpesviruses, suggesting a common mechanism of regulation. We tested one such mechanism, namely how interferon-stimulation influences circRNA expression. 67 circRNAs were upregulated by either IFN-β or -γ treatment, with half of these also upregulated during lytic infection. Using gain and loss of function studies we found an interferon-stimulated circRNA, circRELL1, inhibited lytic HSV-1 infection. We have previously reported circRELL1 inhibits lytic KSHV infection, suggesting a pan-herpesvirus antiviral activity. We propose a two-pronged model in which interferon-stimulated genes may encode both mRNA and circRNA with antiviral activity. This is critical in cases of host shutoff, such as alpha- and gamma-herpesvirus infection, where the mRNA products are degraded but circRNAs escape.

## INTRODUCTION

Herpesviridae is a family of large, double-stranded DNA viruses with a biphasic life cycle, a lytic (replicative) and latent (quiescent, immune evasive) phase. There are nine species known to infect humans, including the alpha-herpesvirus Herpes Simplex Virus-1 (HSV-1), beta-herpesvirus human cytomegalovirus (HCMV), and gamma-herpesvirus Kaposi sarcoma-associated herpesvirus (KSHV). Herpesviruses are a major public health concern with individuals testing seropositive for at least three of the nine species by adulthood (1-6). Infection is asymptomatic for many individuals but, in cases of immune-compromise—such as transplant recipients, neonates, and those with HIV/AIDS—these viruses have devastating effects. HSV-1 commonly causes recurrent oral and genital lesions, but can also cause herpes keratitis, herpetic whitlow, and encephalitis (7). HCMV is the most common congenital infection in addition to a severe opportunistic infection in transplant recipients and individuals with HIV/AIDS (8). KSHV is the etiological agents of several cancers including Kaposi sarcoma and primary effusion lymphoma (9). Murine gamma-herpesvirus 68 (MHV68) has close genetic homology to KSHV and serves as a tractable animal model for pathogenesis (10, 11). Therapeutic agents capable of clearing these viruses or vaccines, for all human herpesviruses excluding Varicella Zoster Virus, are lacking.

Viruses evolve unique mechanisms to invade hosts, alter cellular pathways, and redirect cellular factors for viral processes. In parallel, the host employs a barrage of proteins and RNA species to combat infection. An emerging class of transcripts, circular RNAs (circRNA), has recently been implicated in this host-pathogen arms race (12-16). CircRNAs are single-stranded RNAs circularized by 5’ to 3’ covalent linkages called back-splice junctions (BSJs).

High-throughput sequencing paired with chimeric transcript analysis enables global circRNA detection and quantification (17). These techniques find circRNAs to be ubiquitously expressed in an array of organisms and tissues (18, 19). CircRNAs are also expressed by viruses, including KSHV, Epstein Barr Virus (EBV), human papillomavirus, Merkel cell polyomavirus, hepatitis B virus, and respiratory syncytial virus (20-26). The mechanism underlying host circRNA synthesis, back-splicing, is catalyzed by the spliceosome and regulated by RNA binding proteins (RBPs) and tandem repeat elements which mediate interaction of BSJ flanking sequences (27-30). CircRNAs function as miRNA sponges, protein scaffolds, and transcriptional enhancers (31). CircRNAs are generally classified as noncoding RNAs (ncRNAs), although they possess the capacity for cap-independent translation (32-34). Recently, we identified a host circRNA, circRELL1, that increased the growth of KSHV-infected cells while suppressing the lytic cycle, thereby promoting the viral latency program (16). Additional host circRNAs modulate viral infection (circHIPK3-KSHV, circPSD3-Hepatitis C virus) and are implicated in virus-driven tumorigenesis (circARFGEF1-KSHV, circNBEA-Hepatitis B virus) (14, 15, 35, 36). Furthermore, circRNAs made by spliceosome-independent mechanisms leads to activation of the pattern recognition receptor (PRR), RIG-I (12). Another report found that circRNAs, as a class, sequester the RBP encoded by interleukin enhancer-binding factor 3 (NF90/NF110) and this axis modulates vesicular stomatitis virus infection (13).

As circRNAs lack ends, they are generally resistant to exoribonucleases with approximately 2.5-fold longer half-lives than their linear counterparts (37, 38). CircRNAs are also more stable in the extracellular space, a feature which has led to much interest in their potential use as a diagnostic biomarker (39). Circularity, however, does not prevent susceptibility of circRNAs to endoribonucleases (endoRNases) such as RNase L and RNase P (40, 41). Herpesviruses also express endoRNases, e.g. HSV-1 virion host shutoff (vhs), EBV BamHI fragment G leftward open reading frame 5 (BGLF5), KSHV shutoff and exonuclease (SOX), and MHV68 murine SOX (muSOX). These viral proteins drive a phenomenon called “host shutoff”, which, in part, ablates the immune response by degrading interferon-stimulated genes (42, 43). The viral endoRNases display broad nucleolytic activity *in vitro* relying on viral and host protein adapters *in vivo* to fine tune their RNA substrates (44). These adaptors facilitate a preference for translationally competent RNA leaving ncRNA enriched in the escapee population (45-50). Circularity itself provides some protection from vhs cleavage *in vitro*, however circRNAs containing an internal ribosome entry site can still be targeted (47). In the context of HSV-1 infection, a recent study reported enrichment of circRNAs relative to their colinear gene products, which was not observed in the context of a vhs-null virus (51).

To define host circRNAs commonly regulated by herpesviruses, we performed comparative circRNA expression profiling of cells infected with alpha- (HSV-1), beta- (HCMV), and gamma-herpesviruses (KSHV; MHV68, a murine model of KSHV). We profiled cell culture and animal models, spanning lytic and latent infection. During lytic HSV-1, KSHV, and MHV68 infection, circRNAs were, as a population, unaffected by host shutoff. Ectopic expression assays with human and murine herpesvirus endoRNases confirmed this observation. This agrees with prior reports regarding HSV-1 vhs-mediated decay (47, 51) and expands the observation to gamma-herpesvirus endoRNases. We identified four human and twelve murine circRNAs commonly upregulated after infection across subfamilies of herpesviruses. The most upregulated pathways in our models of HSV-1, HCMV, and KSHV lytic infection were related to immunity.

Thus, we examined if circRNA expression was affected by treatment with various immune stimuli (LPS, poly I:C, CpG) or type I and II interferons (IFN). 67 circRNAs were upregulated by IFN treatment, with half of these also upregulated during viral infection. Finally, we tested if one of these interferon-stimulated circRNAs, circRELL1, echoed the antiviral function of its colinear gene product. Using gain and loss of function studies, circRELL1 was found to inhibit lytic HSV-1 infection. These results echo our prior finding, that circRELL1 inhibits lytic KSHV infection (16), and hints at a common mechanism of action that spans disparate cell types (fibroblast vs. endothelial) and viruses (alpha vs. gamma-herpesviruses). Our data suggests this class of host shutoff escapees may have largely unprobed potential as immunologic effectors.

## RESULTS

### CircRNA profiling of alpha-, beta-, and gamma-herpesvirus infection

As alternative splicing products, circRNAs share almost complete sequence identity with their linear counterparts derived from the same gene. We used CIRCExplorer3-CLEAR to quantify the unique sequence of circRNAs, namely the 5’ to 3’ back splice junctions (52). CIRCExplorer3 also calculates CIRCscore, the number of reads spanning circRNA BSJs (circ fragments per billion mapped bases, circFPB) against reads spanning mRNA forward splice junctions (linear fragments per billion mapped bases, linearFPB). We profiled RNA-Seq data from alpha-, beta-, and gamma-herpesvirus infection, in cell culture and animal models (Fig. 1A, Sup. Fig. 1-1A, Sup. Fig. 1-2A). The data was a combination of our own RNA-Seq data (HSV-1, KSHV, MHV68) in addition to a previously published dataset (HCMV) from Oberstein & Shenk (2017). For all RNA-Seq, excluding the previously published HCMV dataset (53), ERCC (External RNA Controls Consortium) spike-ins were used to control for global transcriptomic shifts caused by infection. We have summarized all the circRNA and gene expression profiling as interactive data tables (Supporting Datasets 1-6) with their utility demonstrated in Sup. Fig. 1-3.

**Figure 1.**
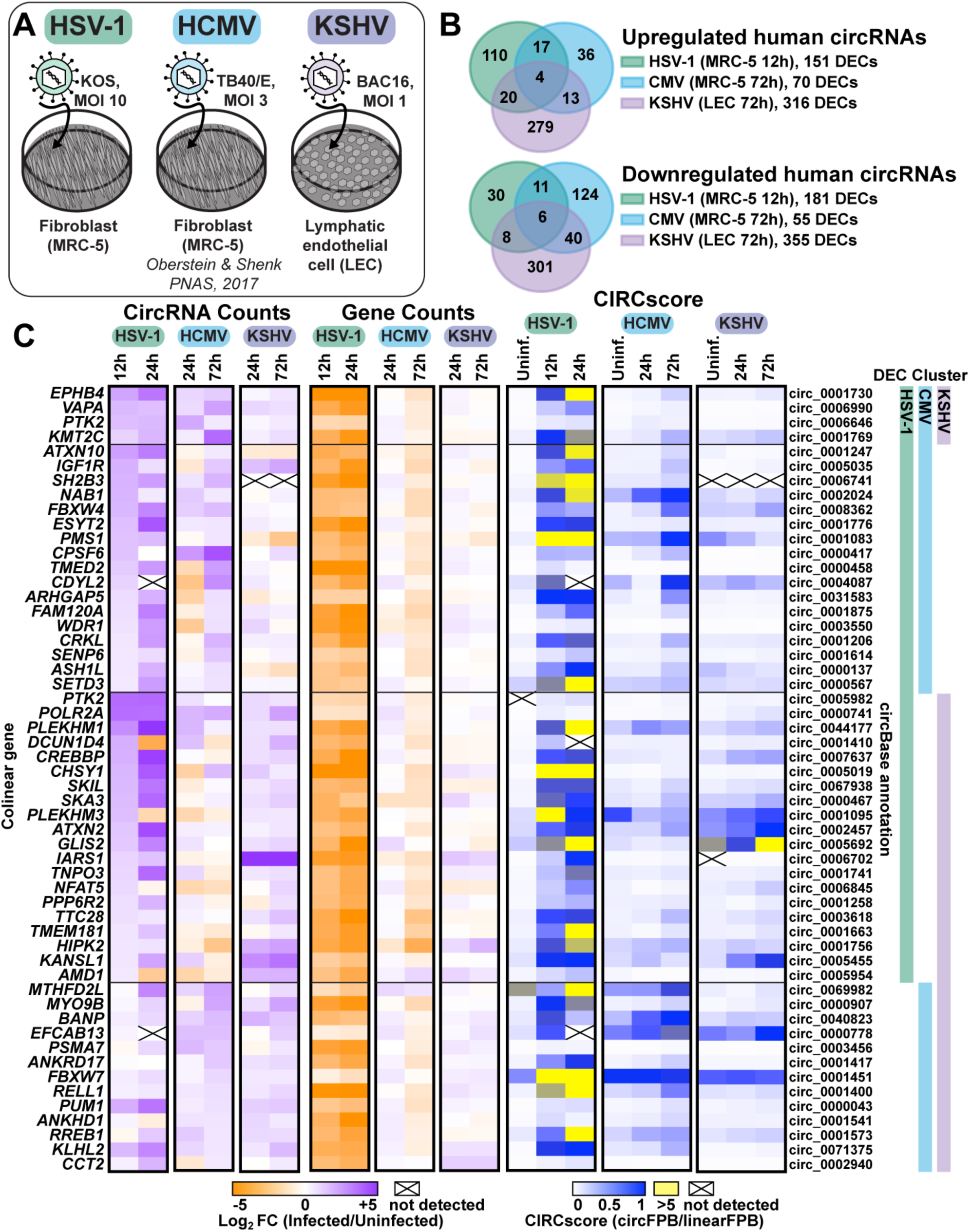
Human circRNAs upregulated in *de novo* lytic infection models. A) Infographic for infection models used in this study. For HSV-1, fibroblasts (MRC-5) were infected with strain KOS at a multiplicity of infection (MOI) of 10 for 12 hours. For HCMV, fibroblasts (MRC-5) were infected with strain TB40/E at an MOI of 3 for 72 hours (Oberstein & Shenk 2017). For KSHV, human dermal lymphatic endothelial cells (LEC) were infected with strain BAC16 at an MOI of 1 for 72 hours. B) Overlap of differentially expressed circRNAs (DECs) detected by bulk RNA-Seq from HSV-1 (n=2-4), HCMV (n=2), and KSHV (n=2) infection. DECs had a raw back splice junction (BSJ) count across the sample set >10, Log_2_FC (fold change)>0.5 or <-0.5, and rank product p-value <0.05. C) Heatmaps for DECs which overlap between viruses, with DEC clusters indicating which virus the circRNA was found to be significantly upregulated within. Data is plotted as CircRNA counts (Log_2_FC Infected/Uninfected normalized BSJ counts), Gene counts (Log_2_FC Infected/Uninfected normalized gene counts), or CIRCscore (circFPB (fragments per billion mapped bases)/linearFPB). Heatmap values are the average of biological replicates. Log_2_FC is relative to a paired uninfected control.

In primary lytic infections we identified 151, 70, and 316 upregulated (log_2_ fold change (log_2_FC) Infected/Uninfected >0.5) human circRNAs for HSV-1, HCMV, and KSHV, respectively (Fig. 1B-C, Sup. Table 1). Four circRNAs were upregulated across viruses (Fig. 1B). A similar number of circRNAs were downregulated after infection (log_2_FC <0.5), with six overlapping between models (Fig. 1B). In HSV-1 infection, disparate circRNA/mRNA expression changes were particularly evident, resulting in dramatic CIRCscore shifts (Fig. 1C). Multiple loci had CIRCscores >5, indicating the circRNA, rather than the colinear mRNA, was the predominant mature transcript for that gene. An increasing CIRCscore with infection was also present for other lytic infection models (Sup. Fig. 1-2C, Fig. 1C). We extended our analysis to mouse models, including HSV-1 infected trigeminal ganglia and MHV68 infected cell lines. We identified 113 murine circRNAs upregulated by HSV-1 infection, although none overlapped between latency, explant-induced reactivation, and drug-enhanced reactivation (Sup. Fig. 1- 1B-C, Sup. Table 1). There were 72 murine circRNAs upregulated by MHV68 infection, with four overlapping between primary infection and lytic reactivation (Sup. Fig. 1-2B-C, Sup. Table 1). There were 12 circRNAs in common between HSV-1 and MHV68 infection models, of these circMed13l (mmu_circ_0001396) was upregulated in all infection models (Sup. Fig. 1-1 and 1- 2).

Differentially expressed circRNAs (DECs) common across disparate virus and cell models hints at a common mode of induction. To investigate this, we performed overrepresentation analysis (ORA) on the colinear genes of circRNAs expressed in human infection models (Sup. Fig. 1-4). ORA identified enrichment of genes involved in cellular senescence for all fibroblast (MRC-5) models, likely representing that infection is performed in G0 cells. Interestingly, genes within the lysine degradation pathway were enriched after infection of HSV-1, HCMV, and KSHV (Sup. Fig. 1-4). One biological function of circRNA is regulation of mRNA expression through miRNA sponging (31). We performed circRNA-miRNA-mRNA network analysis for circRNA commonly upregulated by herpesvirus infection (circEPHB4, circVAPA, circPTK2, and circKMT2C) (Sup. Fig. 1-5). *In silico* analysis predicted miRNA-mRNA interaction nodes which were enriched for mRNA involved in adaptive immunity including MHC complex assembly, TAP complex binding, and peptide antigen stabilization, suggesting a potential role in antiviral immunity for these commonly upregulated circRNAs.

### Global distribution shifts for mRNA, lncRNA, and circRNA during lytic infection

In HSV-1 infection of MRC-5 (Fig. 1C), circRNA upregulation was at odds with the stark decrease in colinear gene expression. A similar trend was visible, albeit less notable, for other lytic infection models (Sup. Fig. 1-2C, Fig. 1C). This finding led us to question if circRNAs were resistant to the global downregulation of host RNAs which occurs during lytic infection. Using ERCC normalized RNA-Seq datasets we plotted read distribution shifts for HSV-1, KSHV, and MHV68 lytic infection (Fig. 2). We compared expression changes for protein-coding (mRNA) genes, long noncoding RNAs (lncRNAs), and circRNAs. In Figure 2A we observed a sharp decrease in host mRNA levels, with a median Log_2_FC of -4.3 (HSV-1 12 hpi), -2.2 (KSHV 72 hpi), and -1.9 (MHV68 18 hpi). This coincides with high levels of viral gene expression. As has been previously reported (48-50), this effect was partially ablated for lncRNAs with median Log_2_FC of -3.9 (HSV-1 12 hpi), -1.3 (KSHV 72 hpi), and -1.5 (MHV68 18 hpi) (Figure 2B). Strikingly, host circRNAs were globally resistant to HSV-1 shutoff and instead exhibited a general upregulation (Log_2_FC +1.3) by 24 hpi (Figure 2C). During KSHV and MHV68 infection, host circRNA expression changes were tri-modal with a down-regulated, unaffected, and up-regulated subpopulation. However, the bulk of circRNA species were again largely unchanged with median Log_2_FC of +0.2 (KSHV 72 hpi) and -0.2 (MHV68 18 hpi). Our analysis demonstrates that host circRNAs, as a species of RNAs, are resistant to the global downregulation of RNA which occurs during lytic herpesvirus infection.

**Figure 2.**
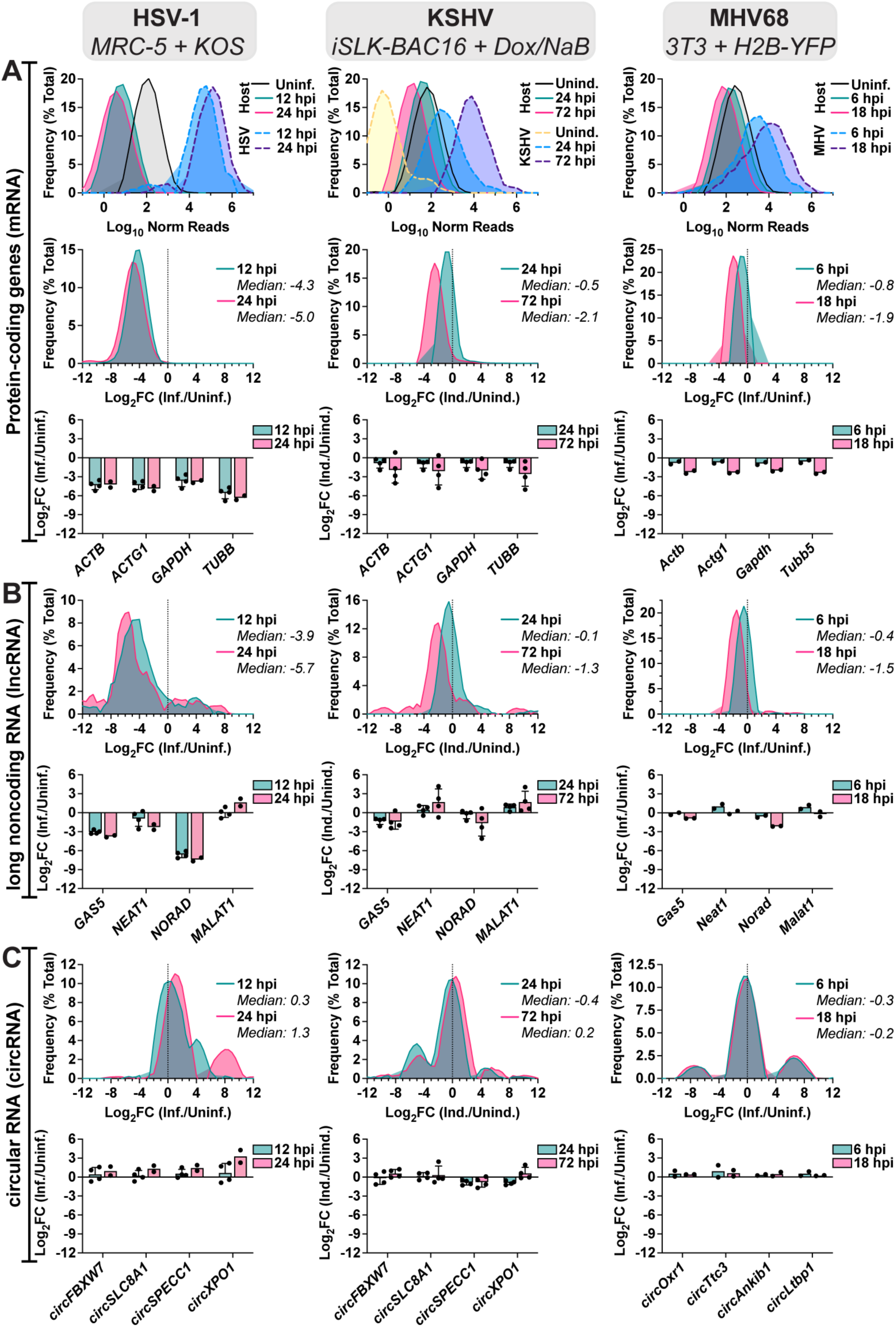
Global distribution shifts for mRNA, lncRNA, and circRNA during lytic infection. Bulk RNA-Seq data from HSV-1 lytic infection (MRC-5 infected with strain KOS MOI 10, n=2- 4), KSHV lytic reactivation (iSLK-BAC16 induced with 1 µg/mL Doxycycline (Dox) 1 mM Sodium Butyrate (NaB), n=4), and MHV68 lytic infection (NIH3T3 infected with strain H2B-YFP MOI 5, n=2). Data is plotted for A) protein-coding genes (top 10,000), B) lncRNAs (top 100), and C) circRNAs (top 100). A-C) Relative frequency distribution was plotted for log_10_ ERCC normalized reads or log_2_FC (Infected/Uninfected or Induced/Uninduced). Log_2_FC for representative genes were plotted, data points are biological replicates, column bars are the average, and error bars are standard deviation.

### CircRNAs are resistant to viral endonuclease mediated decay

Figure 2 plots expression changes across an entire RNA class. To investigate if similar trends occurred for circRNAs and mRNAs derived from the same gene, we evaluated expression shifts for circRNAs (circFPB) and mRNAs (linearFPB) as log_2_FC (Infected/Uninfected) for HSV- 1, KSHV, and MHV68 lytic infection (Fig. 3A). Each dot is a gene that can be alternatively spliced, generating both circRNAs and mRNAs. Echoing our results in Fig. 1C and Fig. 2A, C, circRNA abundance increased while mRNA decreased for vast majority of genes after HSV-1 infection (Fig. 3A). For KSHV and MHV68, mRNA downregulation was consistently more pronounced than circRNA downregulation. Bulk RNA-Seq examines steady-state transcript abundance, averaging the effects of transcriptional activity in addition to co-and post-transcriptional processing such as splicing and decay. To examine what most influences circRNA upregulation we examined our previously published nascent RNA-Seq and ChIP-Seq data (54) for a subset of genes colinear to circRNAs upregulated during HSV-1 infection (Sup. Fig. 3-1). All four genes (*POLR2A, EPHB4, CREBBP, PLEKHM1*) had a drop in RNA Polymerase II (Pol II) and TATA-binding protein (TBP) occupancy by 4 hpi, consistent with published mechanisms of host transcriptional shutoff during HSV-1 infection (55-59). In the case of *EPHB4*, *CREBBP*, and *PLEKHM1* we also observed a drop in 4 thiouridine (4sU)-Seq read coverage, consistent with decreased nascent transcripts. This data suggests circRNA expression changes—for at least this subset—are related to co-or post-transcriptional processing.

**Figure 3.**
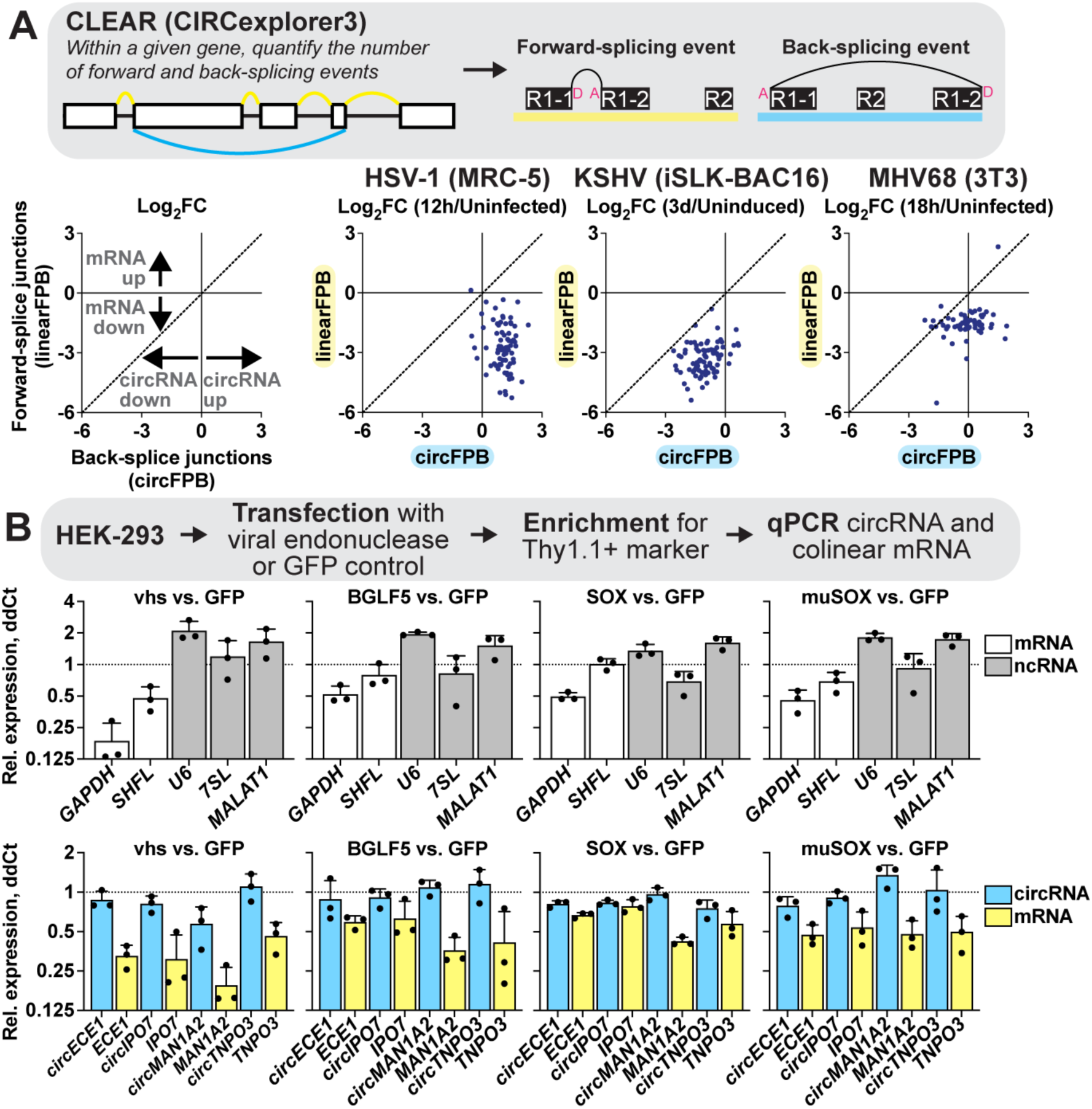
CircRNA are resistant to viral endonuclease mediated decay. A) Bulk RNA-Seq analyzed using CLEAR (CircExplorer3) from HSV-1 (MRC-5 infected with KOS MOI 10 for 12 hours, n=4), KSHV (iSLK-BAC16 reactivated with Dox/NaB for 3 days, n=4), and MHV68 (3T3 infected with H2B-YFP MOI 5 for 18 hours, n=2) infection. Graphs are limited to genes where raw BSJ and forward splice junction (FSJ) counts were >1 across all biological replicates. The average Log_2_FC Infected/Uninfected (HSV-1, MHV68) or Induced/Uninduced (KSHV) was plotted for linearFPB and circFPB, with each dot being a distinct gene. B) HEK-293 cells were transfected for 24 hours with plasmid vectors expressing GFP or viral endonucleases (vhs, BGLF4, SOX, muSOX). RNA was collected and assessed after reverse transcription using qPCR to quantitate mRNAs and noncoding RNAs (ncRNAs). Data is plotted as relative expression (ddCt) using 18S rRNA as the reference gene, and relative to a paired GFP transfected sample. Data points are biological replicates, column bars are the average, and error bars are standard deviation.

Given this observation, we explored if differences in viral endoRNase-mediated decay might explain the disparate expression profiles for circRNA and their colinear genes. Using the approach of Rodriguez *et al.* (2019), plasmids expressing HSV-1 vhs, EBV BGLF5, KSHV SOX, and MHV68 muSOX were transfected into HEK-293 cells. After 24 hours we collected total RNAs and measured transcript levels relative to a GFP vector control (Fig. 3B). As expected, transfection with viral endoRNases decreased *GAPDH* expression (60). Conversely, ncRNAs such as *U6 snRNA*, *7SL*, and *MALAT1* were either unaffected or were increased. We also recapitulated prior findings, regarding escape of the *SHFL* mRNA from SOX-mediated decay (60). We then tested expression changes of circRNAs and colinear mRNAs using divergent or convergent primers, respectively. Across all genes tested and viral endoRNases transfected, circRNA were more resistant to decay as compared to their colinear gene product. Thus, we propose circRNAs as a general class of host shutoff escapees. Our data agrees with prior work from HSV-1 (51), arguing circRNA resistance to viral endoRNases is primarily responsible for the disparate expression changes of circRNAs relative to their colinear mRNA gene products.

### Detection of interferon-stimulated circRNAs (ISCs)

The most significantly upregulated pathways during HSV-1, HCMV, and KSHV infection fell largely within the category of immune responses, with IFN-β and -γ predicted as upstream regulators (Sup. Fig. 4-1). This led us to investigate if circRNA expression may be modulated by innate immune signaling. We treated fibroblast (MRC-5), lymphatic endothelial cell (LEC), and B-cell lymphomas (Akata-, BJAB, Daudi) with immune stimulants including lipopolysaccharide (LPS), CpG DNA, poly I:C, IFN-β and -γ (Sup. Fig. 4-2). A canonical interferon-stimulated gene (ISG), *ISG15*, was measured as a surrogate for immune stimulation. We measured circRELL1 expression, as this circRNA was upregulated in HCMV, KSHV, and to a lesser extent, HSV-1 infection (Fig. 1C). circRELL1 was upregulated in LEC treated with LPS or IFN-γ and B-cell lymphomas treated with poly I:C, LPS, or IFN-β (Sup. Fig. 4-2). *ISG15* activation did not always correlate with circRELL1 expression—notably CpG and poly I:C treatment largely failed to induce expression. Additionally, IFN-β and -γ caused inverse phenotypes in circRELL1 expression when comparing LEC and B-cell models (Sup. Fig. 4-2B). These findings demonstrate that circRELL1 can be upregulated by Toll-like receptor engagement and type I and II interferon stimulation, and the expression profile varies by cell type and mechanism of immune stimulation.

To globally profile interferon-stimulated circRNAs (ISCs) we performed RNA-Seq on fibroblast, lymphatic endothelial, and B-cells treated with IFN-β and -γ. B-cells (Akata) were only treated with IFN-β as we found them refractory to IFN-γ (Sup. Fig. 4-2B). Transcriptomic analysis identified strong upregulation of many canonical ISGs including *OAS1*, *OAS2*, *OASL*, *IFIT1*, *IFITM1*, *MZ1*, *HLA*-*DRB1*, *HLA*-*DQA1*, and *HLA*-*DMA* after IFN treatment (Fig 4A, Supporting Dataset 6). The extent and range of ISGs was most pronounced for B-cells treated with IFN-β (n=4173 DEGs) (Fig. 4A-B) and smallest for fibroblast treated with IFN-β (n=472 DEGs). We identified approximately a dozen interferon-stimulated circRNAs in each model (Fig 4C). Of these, circEPSTI1 (hsa_circ_0000479) was upregulated in all conditions (Fig 4D). We tested if circRELL1 and circEPSTI1 expression changes could be recapitulated in peripheral blood mononuclear cells (PBMCs) and observed a 1.5 and 9-fold increase, respectively, after IFN-β treatment (Sup. Fig. 4-3).

**Figure 4.**
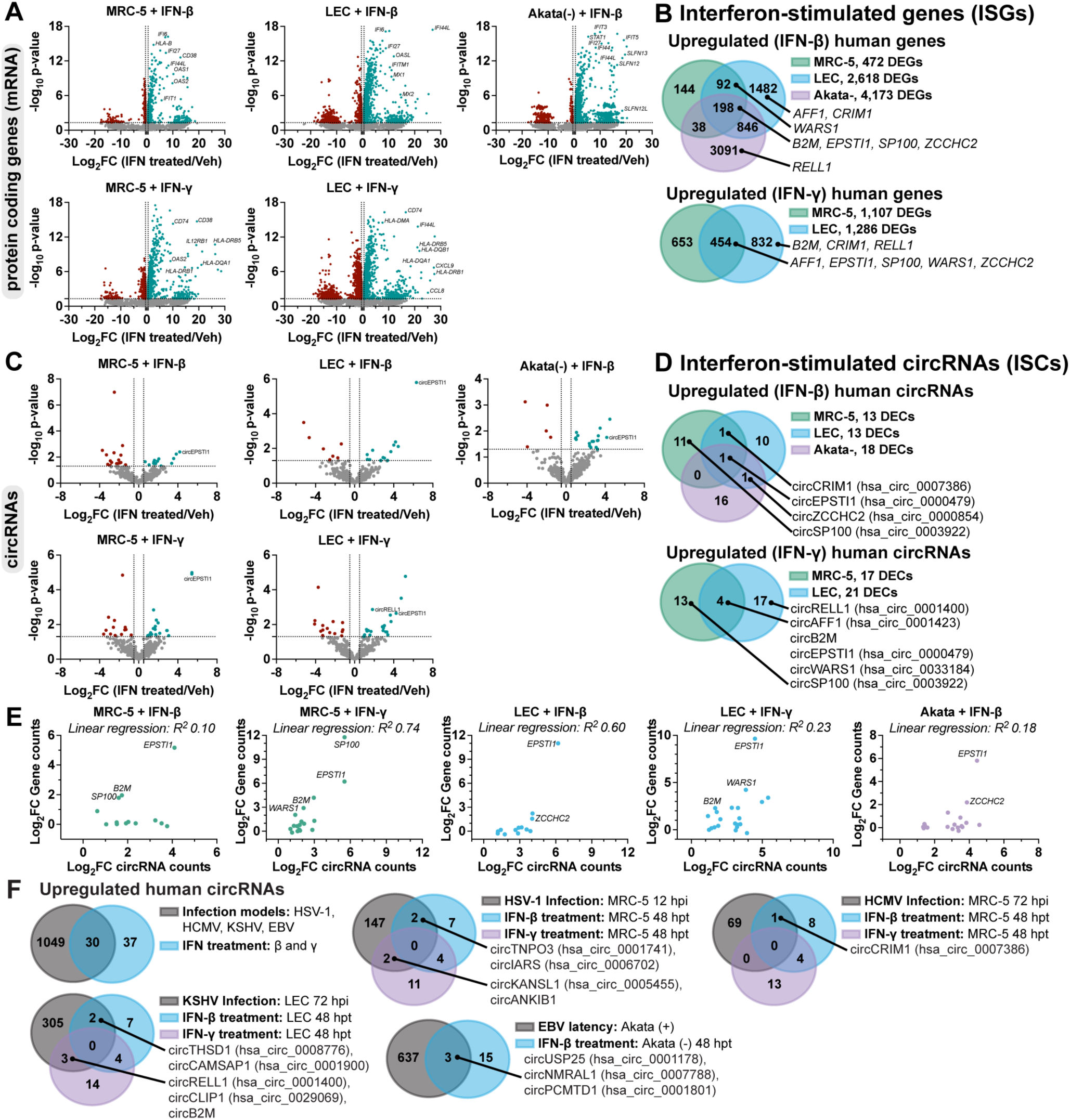
Detection of interferon-stimulated circRNAs (ISCs). A-E) MRC-5, LEC, or Akata-cells were treated with recombinant interferons for 48 hours (n=3). MRC-5 and LEC were treated with IFN-β and -γ (25 ng/mL conc). Akata-were treated with IFN-β (10 ng/mL). Bulk RNA-Seq was performed and data was normalized relative to ERCC spike-in controls. A, C) Volcano plots for ERCC normalized mRNA or circRNA reads. P-values were calculated using EdgeR. Log_2_FC was calculated relative to a paired untreated sample. B, D) Venn diagrams of significantly upregulated circRNA or mRNAs. DECs had a raw BSJ count across the sample set >10, Log_2_FC>0.5 and EdgeR p-value <0.05. DEGs had a Log_2_FC>0.5 and EdgeR p-value <0.05. E) Log_2_FC (stimulated/untreated) was plotted for DECs in B and D relative to their colinear gene reads. R^2^ values are from linear regression analysis. F) Overlap of circRNAs upregulated during herpesvirus infection (Fig. 1) or interferon-stimulation (Fig 4B, D).

A number of colinear mRNAs and circRNAs were stimulated by interferon treatment, including transcripts derived from *CRIM1*, *EPSTI1*, *ZCCHC2*, *SP100*, *AFF1*, *B2M*, and *WARS* (Fig 4B, D). The typical mechanism of ISG induction relies upon promoter activation by factors like IRF3, IRF7, IRF9, STAT1, and STAT2. If this mechanism is similarly responsible for circRNA upregulation, we would expect circRNA levels to correlate with changes in their colinear gene products. We plotted gene expression relationship for ISCs and colinear transcripts (Fig 4E). A subset of ISC-hosting genes, *EPSTI1*, *SP100*, *B2M*, and *WARS*, had a direct relationship between circRNA and colinear gene expression changes. However, this was not globally the case, with linear regression R^2^ values ranging from <0.3 (MRC-5 + IFN-β, LEC + IFN-γ, Akata + IFN-β) to ≥0.6 (MRC-5 + IFN-γ, LEC + IFN-β). We next examined overlaps between our interferon-stimulated and infection-stimulated circRNAs (Fig. 1, Supporting Datasets 1, 2, 3, and 6). Approximately half of the ISCs detected were also upregulated during models of HSV- 1, HCMV, KSHV, or EBV infection (Fig. 4F). These findings demonstrate tunability in circRNA expression with dependence on cell type and immune stimulation.

### circRELL1 restricts HSV-1 lytic infection

In this study, we found circRELL1 to be induced by HSV-1 (1.3-fold), HCMV (1.4-fold), and KSHV (2-fold) infection (Fig. 1C). circRELL1 was also upregulated by immune stimulants including LPS, poly I:C, IFN-β and -γ (Sup. Fig. 4-2 and 4-3). Our lab has previously reported inhibition of lytic KSHV infection by circRELL1 (16). Thus, we questioned if it could be broadly antiviral, by perturbing circRELL1 within the context of HSV-1 infection. To test the impact of loss of function, we depleted circRELL1 48 hours prior to infection using siRNAs targeting the BSJ. MRC-5 cells were infected at a low (0.1 plaque forming units (PFU)/cell) and high (10 PFU/cell) multiplicity of infection (MOI). We achieved significant depletion (4- to 10-fold decrease) of circRELL1 as compared to a Non-Targeting Control (Fig. 5A, D). Viral gene expression for immediate early (IE), early (E), and true late (L2) genes trended upwards, but was largely unaffected by circRELL1 knockdown. Viral entry was likely unaffected as viral genomes at 2 hpi were comparable between Non-Targeting Control and siRNA against circRELL1 (Fig. 5B, E). By 12 hpi, HSV-1 genome copies/cell trended upwards with circRELL1 depletion. The effect of circRELL1 depletion was most apparent when measuring infectious viral yield, with a 1.8- and 2.2-fold increase for low and high MOI infections, respectively (Fig. 5C, F). For gain of function studies, we transduced circRELL1 in MRC-5 with a replication-null lentivirus for 48 hours followed by infection with HSV-1 at high MOI (10 PFU/cell). By 60 hours post-lentivirus infection, circRELL1 expression was 100-fold greater than our lentivirus control that harbors circGFP (Fig. 5G). While we observed no alterations in IE, E, or L2 viral transcripts, viral yield was decreased 4.4-fold by circRELL1 overexpression (Fig. 5F-G). Our loss and gain of function models agree, supporting an anti-lytic role for circRELL1 in HSV-1 infection that appeared to be independent of viral gene expression.

**Figure 5.**
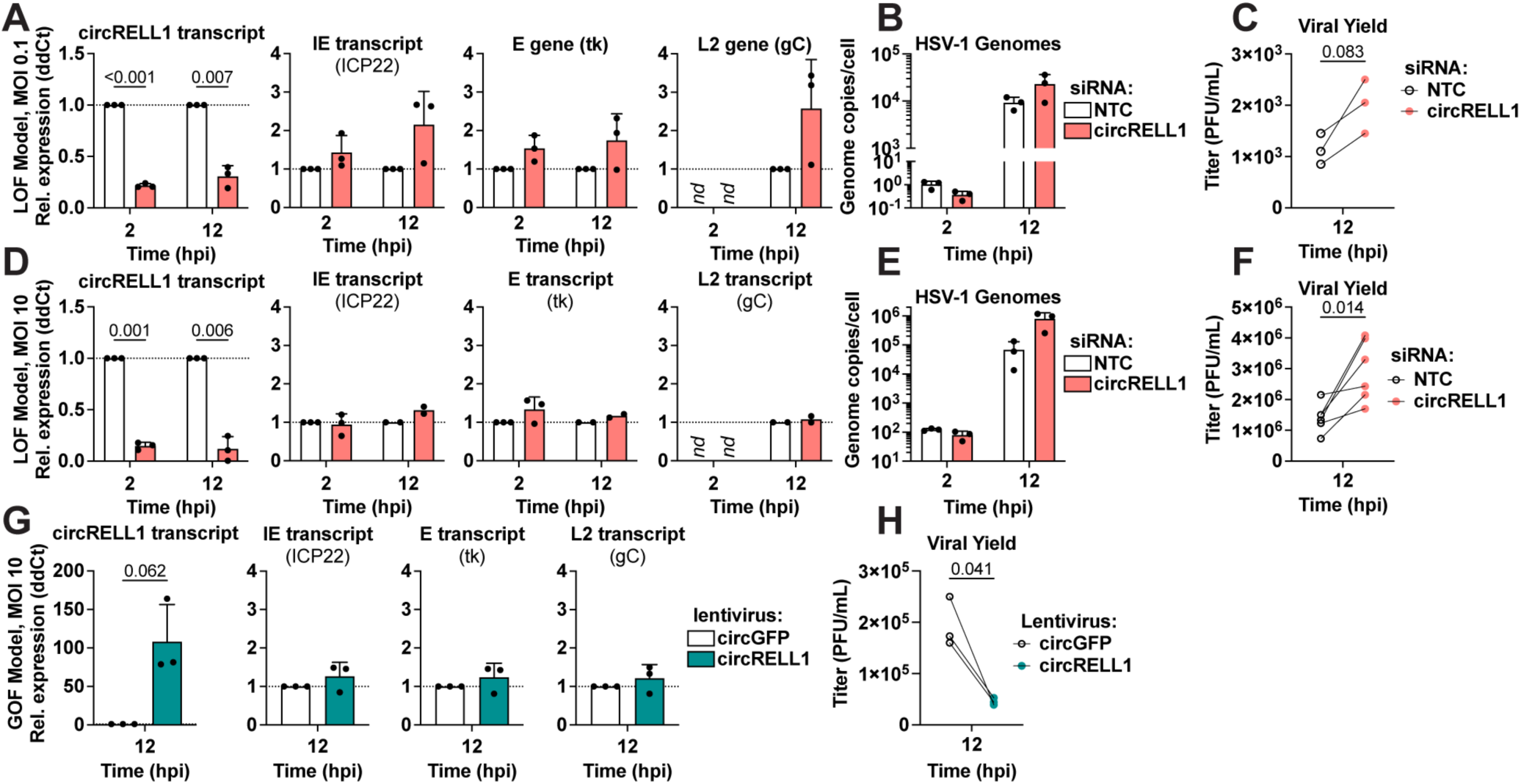
circRELL1 (hsa_circ_0001400) restricts HSV-1 lytic infection. A-F) circRELL1 was depleted in MRC-5 cells using siRNAs for 48 hours and subsequently infected with HSV-1 strain KOS at MOI of 0.1 or 10 PFU/cell. Data is relative to a paired Non-Targeting Control siRNA (NTC). G-H) MRC-5 cells were infected with a lentivirus expressing circRELL1 for 48 hours and subsequently infected with HSV-1 strain KOS at MOI of 10 for 12 hours. Data is relative to a control lentivirus expressing circGFP. A, D, G) RNA was collected from the cell fraction and reverse transcribed. qPCR data is plotted as relative expression (ddCt) using 18S rRNA as the reference gene. B, E) DNA was isolated from the cell fraction and assessed by qPCR for viral and host genome copies. C, F, H) Supernatant was collected at 12 hpi and assessed by plaque assay. Data points are biological replicates, column bars are the average, and error bars are standard deviation. Paired two-tailed t-tests were performed and any p-value <0.1 are shown.

## DISCUSSION

CircRNAs are gaining traction as important factors at the virus-host interface. CircRNAs have an interesting combination of physical features: versatility to interact with DNA, RNA, and proteins simultaneously, longer half-lives than colinear mRNAs, and secretion to extracellular spaces. Some of these features are shared by other molecules like lncRNAs or microRNAs, but circRNAs uniquely possess all those flexible and multi-faceted characteristics. In this study, we probed the role of circRNAs during herpesvirus infection. We performed comparative circRNA expression profiling and identified four host circRNAs commonly upregulated by all subfamilies of human herpesviruses (Fig. 1). Whether there are universal functions of these commonly regulated circRNAs is not clear but over-representation analysis of colinear transcripts showed their enrichment in cell division/senescence pathways and lysine degradation (Sup. Fig. 1-4). *In silico* circRNA-miRNA-mRNA interaction networks predict circEPHB4, circVAPA, circPTK2, and circKMT2C may regulate cellular immunity via repression of miRNA mediated decay of mRNAs involved in antigen presentation (Sup Fig. 1-5). We identified a subset of circRNAs induced by herpesvirus infection and interferon stimulation (Fig. 4). Since circRNAs and mRNAs expressed from the same loci can have distinct targets and functions, they can be thought of as polycistronic genes. We propose a two-pronged model in which interferon-stimulated genes may encode both mRNA and circRNA with immune-regulatory activity (Fig. 6). This is critical in cases of host shutoff, such as alpha-and gamma-herpesvirus infection, where the mRNA product is degraded but circRNA escapes.

**Figure 6.**
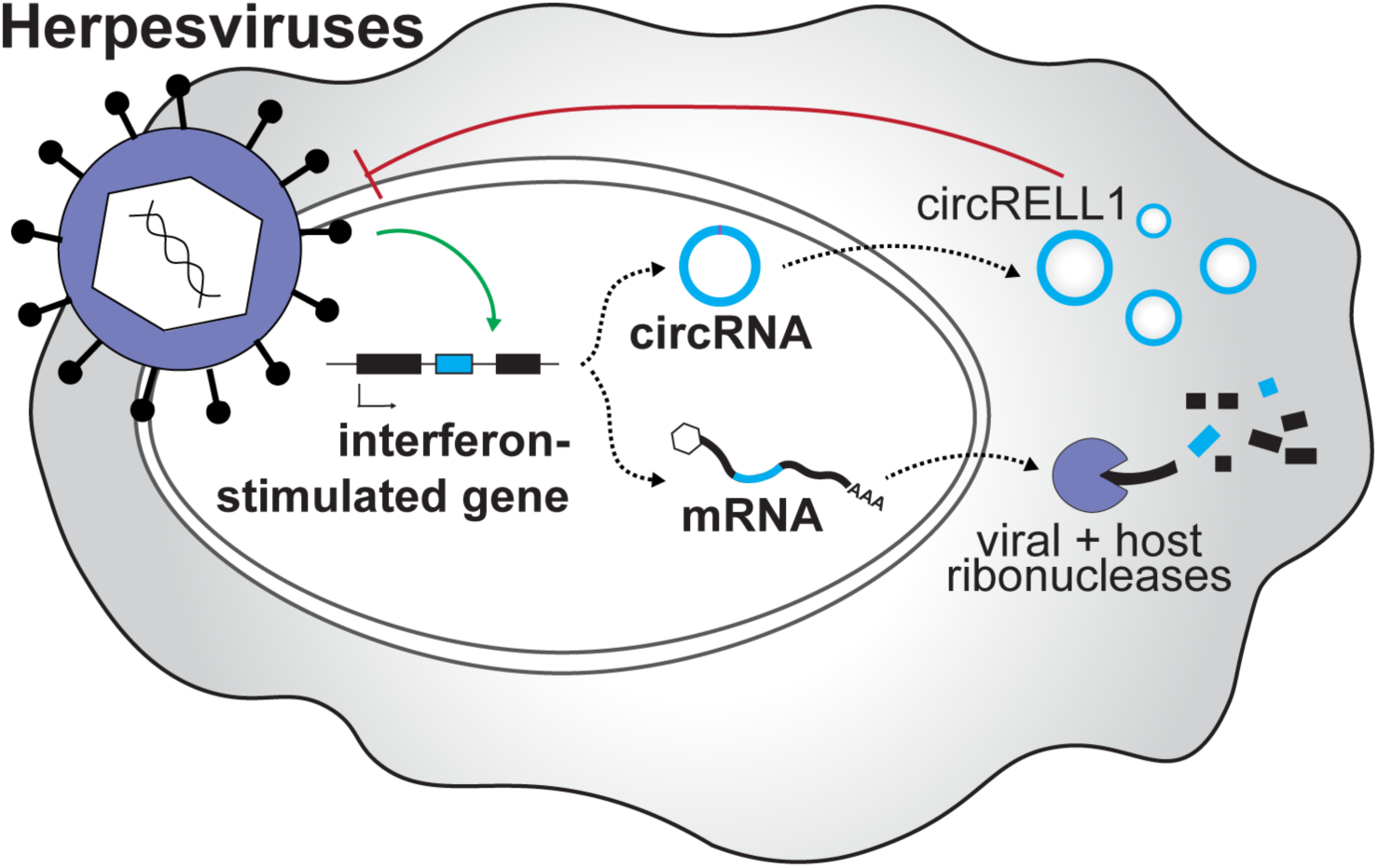
Proposed polycistronic model for interferon-stimulated genes. We propose a polycistronic model in which interferon-stimulated genes can produce both mRNA and circRNA with antiviral activity. This is critical in cases of host shut off, such as alpha-and gamma-herpesvirus infection, where the mRNA product is degraded but circRNA escapes. The interferon-stimulated circRNA, circRELL1, exemplifies this model. EBV, KSHV, and HCMV infection upregulates circRELL1 expression which functions to suppress lytic infection of HSV-1 and KSHV.

CircRNAs were resistant to ectopic expression of HSV-1, EBV, KSHV, and MHV68 endoribonucleases (Fig. 3B), suggesting them to be a general class of host shutoff escapees. These results were echoed during lytic infection, with global circRNA levels generally unaffected as lytic infection progressed (Figure 2C). For the gamma-herpesviruses, KSHV and MHV68, we observed a subpopulation of circRNA which were downregulated by infection. This created a tri-modal circRNA distribution which was not observed for HSV-1. The different circRNA expression modalities for alpha-versus gamma-herpesviruses likely reflects the mechanisms by which specific endoRNases are recruited. HSV-1 vhs targets the 5’ untranslated region (UTR) of mRNAs near the 5’ cap and binding is mediated by the translation factors eIF4H and eIF4AI/II (45-47, 61). CircRNAs may escape vhs degradation as they do not have a 5’ cap or UTR. This mechanism would not apply to gamma-herpesvirus endoRNases (BGLF5, SOX, muSOX) which bind to a degenerate motif with little preference for relative position within the transcript (43, 62). An RNA motif called the SOX resistant element (SRE) can confer resistance to cleavage in a cis-acting, dominant fashion (63, 64). CircRNAs tend to shape characteristic stem structures which may potentially act as SREs (40). RNA modifications or general accessibility may also confer endoRNase resistance. The host is known to use m6A modifications to distinguish self and non-self circRNA (65). Recently, a m6A reader *YTHDC2* was found to be essential for the host shutoff resistance from KSHV SOX (66). RNA Binding Proteins (RBPs) may also play a role in circRNA decay as they can aggregate and cover up to 100% of circRNAs (67), potentially restricting endoRNase substrate recognition. A combination of circRNA sequence and structure likely contributes to the subsets of gamma-herpesvirus endoRNase susceptibility.

CircRNA upregulation in disparate virus and cell models hints at a common mode of induction. As a first line of the antiviral response, the host senses and responds to pathogens through pattern recognition receptors to induce interferons and interferon-stimulated genes. In line with this, the most significantly upregulated pathways during lytic infection were immune-related with IFN-β and -γ predicted as upstream regulators (Sup. Fig. 4-1). We profiled expression after type I and II interferon treatment and identified 67 upregulated circRNAs (Fig. 4), with half of these also upregulated in our infection models. mRNA and circRNA products for several genes including *EPSTI1*, *B2M*, and *ZCCHC2* were induced comparably after interferon stimulation. These interferon-stimulated circRNAs are therefore likely regulated at the level of gene expression. ISCs with disparate expression changes to their colinear gene product (Fig. 4E), may be the results of secondary effects such as altered back-splicing or decay. Zinc finger antiviral protein (ZAP) is upregulated upon IFN stimulation and recruits de-capping enzyme Dcp1 and deadenylase PARN to degrade mRNAs (43). circRNAs are devoid of cap structures or accessible poly(A) tails and likely to resistant to ZAP-mediated degradation, which may result in the accumulation of circRNAs compared to co-linear mRNAs upon IFN stimulation.

An interferon-stimulated circRNA, circRELL1, exemplifies the functionally conserved circRNA. EBV, KSHV, and HCMV infection upregulates circRELL1 expression. We previously showed that circRELL1 has an anti-lytic cycle role during infection with a gamma-herpesvirus, KSHV (16, 21). Here, comparable defects in viral genome replication and infectious progeny after circRELL1 perturbation were observed for the alpha-herpesvirus, HSV-1 (Fig. 5). This functional conservation signifies the importance of commonly regulated circRNAs identified in this study. The mechanism by which this circRNA represses lytic infection is not fully understood. circRELL1 was found to interact with *TTI1* mRNA, a component of mTOR complex, which is targeted by EBV, KSHV, HCMV, and HSV-1 to regulate viral replication (68). Effects of infection-induced circRNAs on mTOR signaling pathway and its consequences thus warrant further study.

## ACKNOWLEDGEMENTS

This work was supported by the Intramural Research Program of the National Cancer Institute (J.M.Z.; 1ZIABC011176, L.T.K; 1ZIABC011953) and National Institute of Allergy and Infectious Diseases (T.M.K.; 1ZIAAI000712). T.T. was funded by a research fellowship from the Japan Society for the Promotion of Science. The funders had no role in study design, data collection and analysis, decision to publish, or preparation of the manuscript. We thank the Center for Cancer Research Sequencing Facility (Frederick, MD) for their assistance in sequencing libraries. The resources of the NIH High-Performance Computing BIOWULF Cluster were utilized for all our computational needs. We thank Mandy Muller (University of Massachusetts Amherst) for plasmids containing viral endonucleases. We thank Neal DeLuca (University of Pittsburgh) for HSV-1 (strain KOS). We thank Bill Sugden (University of Wisconsin Madison) for providing B cell lymphoma cell lines.

## AUTHOR CONTRIBUTIONS

SD, TT, JA, TK, and LK designed and performed the experiments. SD and VK analyzed deep-sequencing data. SD, TT, and JZ interpreted data and wrote the manuscript. SD and TT contributed equally to this research. All authors contributed to the article and approved the submitted version.

## METHODS

### Cells and Viruses

Vero (ATCC #CCL-81) were maintained in Dulbecco’s modified eagle medium (DMEM, Gibco #11965-092) supplemented with 5% fetal bovine serum (FBS, Gibco #16000044), 1 mM sodium pyruvate (Gibco # 11360070), 2 mM L-glutamine (Gibco # A2916801), 100 U/mL penicillin-streptomycin (Gibco # 15070063). MRC-5 (ATCC #CCL-171) were maintained in DMEM (Gibco) supplemented with 10% FBS, 1 mM sodium pyruvate, 2 mM L-glutamine, 100 U/mL penicillin-streptomycin. NIH 3T3 (ATCC #CRL-1658) and NIH 3T12 (ATCC #CCL-164) were maintained in DMEM (Corning #10-017-CV) supplemented with 8% FBS, 2 mM L-glutamine, 100 U/mL penicillin-streptomycin. A20 HE-RIT B cells harbor a recombinant MHV68 expressing a hygromycin-eGFP cassette (69) and doxycycline-inducible RTA (70). A20 HE-RIT were maintained in RPMI supplemented with 10% FBS, 100 U/mL penicillin-streptomycin, 2 mM L-glutamine, 50 µM beta-mercaptoethanol, 300 µg/ml hygromycin B, 300 µg/mL gentamicin, and 2 µg/mL puromycin. iSLK-BAC16 (71) were maintained in DMEM (Gibco #11965-092) supplemented with 10% Tet system-approved FBS (Takara #631368), 50 µg/mL hygromycin B, 0.1 mg/mL gentamicin, 1 µg/mL puromycin, 100 U/mL penicillin-streptomycin. 293T (ATCC #CRL-3216) cells were maintained in DMEM supplemented with 10% FBS and 100 U/mL penicillin-streptomycin. Lymphoma cell lines, EBV-positive and negative Akata (designated with (+) or (-)) (72), Daudi (ATCC #CCL-213), and BJAB (DSMZ *#*ACC757) were maintained in Roswell Park Memorial Institute (RPMI) 1640 (Gibco #11875093) supplemented with 10% FBS and 100U/ml penicillin-streptomycin. HDLEC (PromoCell #C-12216) were maintained in EBM-2 basal medium (Lonza #CC-3156) supplemented with EGM-2 SingleQuots supplements (Lonza #CC-4176).

### Virus Stock Preparation and Titration

#### HSV-1

Vero cells were infected with KOS (73) or strain 17 at a low multiplicity of infection (MOI; ∼0.01 plaque forming units (PFU)/cell) and harvested when cells were sloughing from the sides of the vessel. Supernatant and cell fraction were collected and centrifuged at 4,000 x g 4°C for 10 minutes. The subsequent supernatant fraction was reserved. The pellet fraction was freeze (-80°C 20 min)/thawed (37°C 5 min) for three cycles, sonicated for 1 minute, and centrifuged at 2,000 x g 4°C for 10 minutes. The final virus stock was composed of the cell-associated virus and reserved supernatant virus fractions.

#### KSHV

iSLK-BAC16 cells were induced with 1 µg/mL doxycycline and 1 mM sodium butyrate for 3 days. Cell debris was removed from the supernatant fraction by centrifuging at 2,000 x g 4°C for 10 minutes and filtering with a 0.45 polyethersulfone membrane. Virus was concentrated after a 16,000 x g 4°C 24 hour spin and resuspended in a low volume of DMEM media (approx. 1000-fold concentration). To assess viral infectivity, LECs were infected with serial dilutions of BAC16 stock and assessed using CytoFlex S (Beckman Coulter) for GFP+ cells at 3 days post infection. BAC16 contains a constitutively expressed GFP gene within the viral genome. Based on these assays, BAC16 stock was used at a 1:60 dilution, resulting in 70% infection for LEC (MOI 1).

#### MHV68

NIH 3T12 based cell lines were infected at a low MOI with MHV68 until 50% cytopathic effect was observed. Infected cells and conditioned media were dounce homogenized and clarified at 600 x g 4°C for 10 minutes. Clarified supernatant was further centrifuged at 3,000 x g 4°C for 15 min and then 10,000 x g 4°C for 2 hrs to concentrate 40- fold in DMEM. H2B-YFP was prepared and titered using plaque assays in NIH 3T12 cells.

### *De Novo* Infection

#### HSV-1

Confluent MRC-5 cells were infected with 0.1 or 10 PFU per cell. Virus was adsorbed in PBS for 1 hr at room temperature. Viral inoculum was removed, and cells were washed quickly with PBS before adding on DMEM media supplemented with 2% FBS. 0 hour time point was considered after adsorption of infected monolayers when cells were place at 37°C. <u>KSHV.</u> Subconfluent LEC were infected with BAC16 at an approximate MOI of 1 (70% cells infected), as assessed by GFP+ cells at 3 dpi. Virus was adsorbed in a low volume of media for 8 hr at 37°C, after which viral inoculum was removed and replaced with fresh media. 0 hour time point was when virus was added and cells were first placed at 37°C.

#### MHV68

Subconfluent NIH3T3 fibroblasts were infected with 5 PFU per cell. Virus was adsorbed in a low volume of DMEM media supplemented with 8% FBS for 1 hr at 37°C, prior to overlay with fresh media. 0 hour time point was when virus was first added and cells were placed at 37°C.

### HSV-1 Mouse Infections

Female 8-week-old BALB/cAnNTac mice were infected with 10^5^ PFU HSV-1 (strain 17) via the ocular route. Latently infected trigeminal ganglia were harvested approximately four weeks after primary infection and immediately processed. For explant-induced reactivation, latently infected trigeminal ganglia were explanted into culture (DMEM/1% FBS) for 12 hours at 37°C/5% CO_2_ in the presence of vehicle (DMSO) or 2 μM JQ1+ to enhance reactivation (Cayman Chemical CAS: 1268524-70-4). Pools of 6 ganglia were homogenized in 1 ml TriPure isolation reagent (Roche) using lysing matrix D on a FastPrep24 instrument (3 cycles of 40 seconds at 6 m/s). 0.2 ml chloroform was added for phase separation using phase lock gel heavy tubes and RNA isolation from the aqueous phase was obtained by using ISOLATE II RNA Mini Kit (Bioline). RNA quality was verified with Agilent 2100 Bioanalyzer System using RNA Nano Chips (Agilent Technologies). All animal care and handling were done in accordance with the U.S. National Institutes of Health Animal Care and Use Guidelines and as approved by the National Institute of Allergy and Infectious Diseases Animal Care and Use Committee (Protocol LVD40E, T.M.K.).

### Lytic Reactivation

#### KSHV

Subconfluent monolayers of iSLK-BAC16 were induced with 1 µg/mL doxycycline, 1 mM sodium butyrate in DMEM media supplemented with 2% Tet system-approved FBS. 0 hour time point was when induction media was added and cells were first placed at 37°C.

#### MHV68

One day prior to induction, A20 HE-RIT cells were subcultured at a 1:3 dilution in media lacking antibiotics. Cells were seeded subconfluently and induced for 24 hours with RPMI media containing 5 µg/ml doxycycline and 20 ng/ml 12-O-tetradecanoylphorbol-13- acetate (TPA).

### rRNA-depleted total RNA-Seq

Total RNA was isolated from cells using the Direct-zol RNA MiniPrep kit (Zymo Research R2053), following manufacturer’s instructions. ERCC spike-in controls (ThermoFisher 4456740) were added to 500-1000 ng of total RNA. RNA was sent to the NCI CCR-Illumina Sequencing facility for library preparation and sequencing. RNA was rRNA depleted and directional cDNA libraries were generated using either Stranded Total RNA Prep with Ribo-Zero Plus (Illumina # 20040525) or TruSeq Stranded Total RNA Ribo-Zero Gold (Illumina #RS- 122-2303). 2-4 biological replicates were sequenced for all samples. Sequencing was performed at the National Cancer Institute Center for Cancer Research Frederick Sequencing Facility using the Illumina NextSeq 550 or Illumina NovaSeq SP platform to generate 150 bp paired-end reads.

### Oligos

A list of all oligos used is located in Supplementary Table 2.

### RNA extraction and RT-qPCR

Total RNA was extracted with Direct-zol RNA miniprep kit with on-column DNase I digestion (Zymo Research #R2053). 0.5 to 1 μg of total RNA was used for reverse-transcription with ReverTra Ace qPCR RT master mix (Toyobo #FSQ-101) and quantitative PCR (qPCR) was performed with Thunderbird Next SYBR qPCR mix (Toyobo #QPX-201) and StepOnePlus real-time PCR system (ThermoFisher) following manufacturer’s instructions.

### Measuring viral genomes

The cell fraction was isolated from infection models. Cell pellets were washed with 1x PBS and lysed using 0.5% SDS, 400µg/mL proteinase K, 100 mM NaCl. Samples were incubated at 37°C for 12-18 hours and heat inactivated for 30 minutes at 65°C. DNA samples were serial diluted 1:1000 and measured using qPCR with primers specific to HSV-1 UL23 and human *GAPDH*. Standard curves were generated using purified genomic stocks (HSV-1 bacterial artificial chromosome and human genome Promega #G1471). Absolute copy number of genomic stocks was determined using droplet digital PCR (Biorad QX600). Values were plotted as follows: 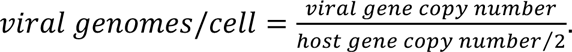

### Viral nuclease ectopic expression

3×10^6^ 293T cells were seeded to 10 cm petri dishes and incubated overnight. Cells were transfected with 8 µg of plasmid DNAs (pCMV-Thy1.1-F2A-dsGFP/muSOX/SOX/vhs/BGLF5) using 48 μl Transporter 5 (Polysciences #26008) and 1ml Opti-MemI (Gibco #31985070). After 24 hours, cells were resuspended in 5 ml staining buffer [1x PBS Gibco # 10010023 supplemented with 2mM ethylenediaminetetraacetic acid (Sigma) and 0.5% FBS (Gibco)] and mouse Thy1.1-expressing cells were magnetically enriched with CD90.1 MicroBeads (Mlitenyi #130-121-273). Cells were stained with Alexa Fluor 647 anti-mouse Thy-1.1 Antibody (BioLegend; clone OX-7) and enrichment was confirmed with CytoFlex S (Beckman Coulter). 70∼80% cells were positive for Thy1.1 after sorting and lysed with TRI reagent (Zymo Research #R2050-1) for RT-qPCR.

### Interferon stimulation

Confluent monolayers of MRC-5 and LEC were treated with 25 ng/mL recombinant human IFN-β and γ (Peprotech #300-02BC and #300-02) in the culture media. For Akata(-) 10 ng/mL IFN-β was added to culture media. After 48 hours RNA was isolated from the cell fraction.

## BIOINFORMATIC ANALYSIS

### Gene quantitation

RNA-Sequencing reads were trimmed using Cutadapt (74) and the following parameters: -- pair-filter=any, --nextseq-trim=2, --trim-n, -n 5, --max-n 0.5, -0 5, -q 20, -m 15. Trimmed reads were mapped using STAR (75) with 2-pass mapping to concatenated genome assemblies which contain the host genome (hg38 or mm39) + virus genome (KT899744.1, NC_001806.2, NC_006273.2, NC_009333.1, MH636806.1) + ERCC spike-in controls. Details on mapping assemblies are included below. RNA STAR mapping parameters are as follows: -- outSJfilterOverhangMin 15 15 15 15, --outFilterType BySJout, --outFilterMultimapNmax 20, -- outFilterScoreMin 1, --outFilterMatchNmin 1, --outFilterMismatchNmax 2, -- outFilterMismatchNoverLmax 0.3, --outFilterIntronMotifs None, --alignIntronMin 20, -- alignIntronMax 2000000, --alignMatesGapMax 2000000, --alignTranscriptsPerReadNmax 20000, --alignSJoverhangMin 15, --alignSJDBoverhangMin 15, --alignEndsProtrude 10 ConcordantPair, --chimSegmentMin 15, --chimScoreMin 15, --chimScoreJunctionNonGTAG 0 --chimJunctionOverhangMin 18, --chimMultimapNmax 10. RNA STAR GeneCount (per gene read counts) files were used for transcript quantitation. The only RNA-Seq data not generated in-house is from HCMV infection (53). This data did not contain ERCC spike-in controls and thus was normalized as Transcripts per Million (TPM). For all other models, ERCC reads were used to generate standard curves similar to (76), using their known relative concentrations. All biological replicates had ERCC derived standard curves with R^2^>0.9. ERCC normalized gene counts were calculated as follows:

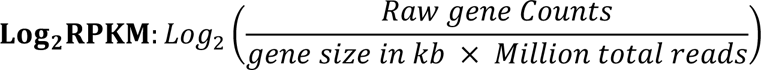

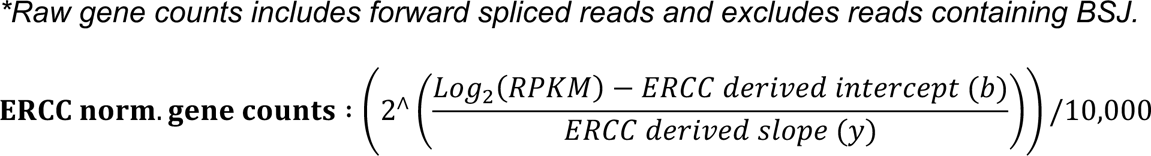

### CircRNA quantitation

RNA-Seq data was trimmed (Cutadapt) and aligned (STAR, 2-pass) as described in the gene quantitation section above. Note that we required a minimum of 18 nucleotides flanking any chimeric BSJ calls to ensure high-confidence in circRNA quantitation. Back-splice junctions (BSJ) were quantified using CIRCExplorer3 (CLEAR) pipeline (52) and normalized as TPM (HCMV data) or relative to ERCC spike in controls (all other data). BSJ variants are reported relative to their colinear gene products and circbase annotations (http://www.circbase.org/) (77). Gene length for circRNA was treated as 0.15 kb as that is the total read length and full circRNA size is unknown. ERCC normalized circRNA counts were calculated as above.

### Differentially expressed circRNA (DEC) calling

#### Bulk RNA-Seq data

Up or down-regulated circRNAs had a raw BSJ count across the sample set >10, Log_2_FC>0.5, and p-value <0.05. With the exception of interferon stimulated RNA-Seq data (Fig. 4), significance was calculated by rank product paired-analysis with RankProd R package (78). For interferon stimulated RNA-Seq data EdgeR was used to calculate statistical significance.

#### EBV microarray data

Data was previously published in (16), comparing Akata(+) and Akata(-) cells assessed by microarray (074301 Arraystar Human CircRNA microarray V2). Upregulated circRNAs had a Log_2_FC>0.5 and p-value <0.05. Significance was calculated by rank product paired-analysis with RankProd R package (78).

### Genome assemblies

<u>HSV-1, strain KOS:</u> KT899744.1, with the corresponding coding sequence (CDS) annotation used for transcript quantification

<u>HSV-1, strain 17:</u> NC_001806.2, with the corresponding CDS annotation used for transcript quantification

<u>HCMV:</u> NC_006273.2, with the corresponding CDS annotation used for transcript quantification

<u>KSHV:</u> NC_009333.1, with the corresponding CDS annotation used for transcript quantification

<u>MHV68:</u> MH636806.1 (79) modified to remove the beta-lactamase gene (Δ103,908-105,091), with the corresponding CDS annotation used for transcript quantification

<u>Human:</u> hg38, gencode.v36

<u>Mouse:</u> mm39, gencode.vM29

<u>ERCC Spike-In:</u> available from ThermoFisher (#4456740)

## DATA AVAILABILITY

Additional information about data analyzed in this study is present in Supplementary Table 3.

<u>HSV-1 infection:</u> SRR19779319, SRR19779318, SRR19787559

<u>HSV-1 murine infection:</u> SRR19792335, SRR19792334, SRR25824398, SRR25824397, SRR25824394, SRR25824396

<u>HCMV lytic infection:</u> SRR5629593, SRR5629594, SRR5629591, SRR5629592, SRR5629589, SRR5629590, SRR5629587, SRR5629588, SRR5629577, SRR5629578, SRR5629575, SRR5629576, SRR5629573, SRR5629574, SRR5629571, SRR5629572

<u>KSHV infection:</u> SRR20020769, SRR20020770, SRR20020761, SRR20020757, SRR20020758, SRR25816558, SRR25816557, SRR25816556

<u>MHV68 infection:</u> SRR19792326, SRR19792325, SRR19792324, SRR19792321, SRR25823338, SRR25823339

<u>Interferon stimulation:</u> SRR25905055, SRR25905049, SRR25905048, SRR25905050, SRR25905054, SRR25905051, SRR25905053, SRR25905052

<u>EBV circRNA microarray data:</u> GSE206824

