## Supplementary information for "Interferon induced circRNAs escape herpesvirus host shutoff and suppress lytic infection"

**This PDF file includes:**

Supplementary methods
Legend for Table S1
Tables S2 to S3
Legends for Datasets S1 to S6
Figures S1-1, S1-2, S1-3, S1-4, S1-5, S 3-1, S 4-1, S4-2, S4-3
SI References

**Other supporting materials for this manuscript include the following:**

Table S1
Datasets S1 to S6

### SUPPLEMENTARY METHODS

#### Immune stimulation of cell culture models

Confluent monolayers of MRC-5 and LEC were treated with a variety of immunostimulants. 10 ug/mL lipopolysaccharide (Millipore Sigma L3024), 4 ug/mL CpG DNA (IDT), 10 ug/mL poly I:C (Millipore Sigma #P1530), or 25 ng/mL recombinant human IFN-β and γ (Peprotech #300-02BC and #300-02) was added to culture media. After the indicated treatment time (8, 24, 48, or 72 hours), RNA was isolated from the cell fraction. For Akata-, Daudi, or BJAB, 10 ug/mL lipopolysaccharide, 10 ug/mL polyI:C, 10 ng/mL IFN-β, or 500 ng/mL IFN-γ were added to culture media. After 24 hours RNA was isolated from the cell fraction.

#### Immune stimulation of PBMCs

Human peripheral blood mononuclear cells (PBMCs) were prepared from buffy coats. Buffy coats from two different donors were purified with Ficoll-Paque PLUS (GE Healthcare #17144003). PBMCs were treated with red blood cell lysis buffer (BioLegend #420301) and washed three times with phosphate-buffered saline (Gibco #10010023). 10 or 30 ng/mL recombinant human IFN-β (Peprotech #300-02BC) was added to the culture media. After 24 hours RNA was isolated from the cell fraction.

#### *In silico* circRNA-miRNA-mRNA network predictions

Four commonly-regulated human circRNAs (hsa_circ_0001730, hsa_circ_0006990, hsa_circ_0006646, hsa_circ_0001769) were examined for miRNA-binding sites using CircInteractome (1). Experimentally supported miRNA-mRNA interactions were then added to the network based on DIANA-TarBase v8.0 (2). Cytoscape 3.10 was used for visualizing the network (3). Overrepresentation analysis of predicted target miRNAs was performed with miEAA 2.0 (4). Enriched Gene Ontology terms (http://geneontology.org), p-values, and miRNAs that were predicted to be regulated by circRNAs are shown.

#### Overrepresentation analysis (ORA)

ORA was performed using a list of genes colinear to circRNAs detected in a given model (>10 raw BSJ read counts). ORA was performed using WebGestalt (5) and plotted using SRplot (https://www.bioinformatics.com.cn/en).

#### Ingenuity pathway analysis (IPA)

IPA (Qiagen) was performed on *de novo* lytic infection RNA-Seq datasets for HSV-1 (MRC-5 infected at MOI (multiplicity of infection) of 10 for 12 hours, 4 biological replicates), HCMV (MRC-5 infected at MOI of 3 for 72 hours, 2 biological replicates), and KSHV (LEC infected at MOI of 1 for 72 hours, 2 biological replicates). $Log_2$ fold change ($Log_2FC$, Infected/Uninfected) was calculated from ERCC normalized data. Only genes with an average $Log_2FC$ >1 were used for IPA. Bubble plots were generated using SRplot (https://www.bioinformatics.com.cn/en).

### SUPPORTING DATASETS

**Supporting Dataset 1. Transcriptomic data from HSV-1 human infection models**
Transcriptomic data from bulk RNA-Seq data for MRC-5 infected with HSV-1 strain KOS at MOI of 10 plaque-forming unit (PFU)/cell for 12 or 24 hours. Data was mapped to a concatenated genome assembly containing hg38 (gencode.v36), KT899744.1, and ERCC (External RNA Controls Consortium) spike-ins. Tab 1-GenePlotter) Query ERCC normalized gene expression data from the samples by inputting gene symbol. Tab 2-CircPlotter) Query ERCC normalized back splice junction (BSJ) data from the samples by inputting circBase annotation. Tab 3-Summary data) RNA Star 2-pass mapping log output file information, including input reads and those mapped to the genome assemblies. Tab 4-Raw counts) CircRNA counts are generated by CircExplorer3 (6) mapping and include only BSJ reads. All other gene counts are outputs from RNA STAR GeneCount (per gene read counts). Tab 5-ERCC norm counts) BSJ or gene counts normalized using ERCC spike-in reads. Tab 6-Log2FC) $Log_2FC$ (Infected/Uninfected) for ERCC normalized counts.

**Supporting Dataset 2. Transcriptomic data from HCMV infection models**
Transcriptomic data from bulk RNA-Seq data for MRC-5 infected with HCMV strain TB40/E at MOI of 3 PFU/cell for 24 or 72 hours (7). Data was mapped to a concatenated genome assembly containing hg38 (gencode.v36) and NC_006273.2. Tab 1-GenePlotter) Query TPM (transcript per million) normalized gene expression data from the samples by inputting gene symbol. Tab 2-CircPlotter) Query TPM normalized BSJ data from the samples by inputting circBase annotation. Tab 3-Summary data) RNA Star 2-pass mapping log output file information, including input reads and those mapped to the genome assemblies. Tab 4-Raw counts) CircRNA counts are generated by CLEAR (CircExplorer3) mapping and include only BSJ reads. All other gene counts are outputs from RNA STAR GeneCount (per gene read counts). Tab 5-TPM norm counts) BSJ or gene counts normalized as TPM. Tab 6-Log2FC) $Log_2FC$ (Infected/Uninfected) for TPM normalized counts.

**Supporting Dataset 3. Transcriptomic data from KSHV infection models**
Transcriptomic data from bulk RNA-Seq data for i. LEC infected with KSHV strain BAC16 at MOI of 1 PFU/cell for 24 or 72 hours or ii. iSLK-BAC16 reactivated with 1 mM Sodium Butyrate 1 ug/mL Doxycycline for 24 or 72 hours. Data was mapped to a concatenated genome assembly containing hg38 (gencode.v36), NC_009333.1, and ERCC spike-ins. Tab 1-GenePlotter) Query ERCC normalized gene expression data from the samples by inputting gene symbol. Tab 2-CircPlotter) Query ERCC normalized BSJ data from the samples by inputting circBase annotation. Tab 3-Summary data) RNA Star 2-pass mapping log output file information, including input reads and those mapped to the genome assemblies. Tab 4-Raw counts) CircRNA counts are generated by CLEAR (CircExplorer3) mapping and include only BSJ reads. All other gene counts are outputs from RNA STAR GeneCount (per gene read counts). Tab 5-ERCC norm counts) BSJ or gene counts normalized using ERCC spike-in reads. Tab 6-Log2FC) $Log_2FC$ (Infected/Uninfected) for ERCC normalized counts.

**Supporting Dataset 4. Transcriptomic data from HSV-1 mouse infection models**
Transcriptomic data from bulk RNA-Seq data for i. mice trigeminal ganglia (TG) latently infected with HSV-1 strain 17 for 4 weeks, or ii. TG explants from mice infected with HSV-1 strain 17 for 5 weeks and grown in the presence or absence of 2 uM JQ1 (Selleckchem, Houston TX) for 12 hours. Data was mapped to a concatenated genome assembly containing mm39 (gencode.vM29), NC_001806.2, and ERCC spike-ins. Tab 1-GenePlotter) Query ERCC normalized gene expression data from the samples by inputting gene symbol. Tab 2-CircPlotter) Query ERCC normalized BSJ data from the samples by inputting circBase annotation. Tab 3-Summary data) RNA Star 2-pass mapping log output file information, including input reads and those mapped to the genome assemblies. Tab 4-Raw counts) CircRNA counts are generated by CLEAR (CircExplorer3) mapping and include only BSJ reads. All other gene counts are outputs from RNA STAR GeneCount (per gene read counts). Tab 5-

ERCC norm counts) BSJ or gene counts normalized using ERCC spike-in reads. Tab 6-Log2FC) $Log_2FC$ (Infected/Uninfected) for ERCC normalized counts.

**Supporting Dataset 5. Transcriptomic data from MHV68 infection models**
Transcriptomic data from bulk RNA-Seq data for (i) NIH 3T3 infected with MHV68 strain H2B-YFP MOI of 5 PFU/cell for 6 or 18 hours, or (ii) A20 HE-RIT reactivated with 5 ug/ml Doxycycline and 20 ng/ml 12-O-tetradecanoylphorbol-13-acetate (TPA) for 6 or 24 hours. Data was mapped to a concatenated genome assembly containing mm39 (gencode.vM29), MH636806.1, and ERCC spike-ins. Tab 1-GenePlotter) Query ERCC normalized gene expression data from the samples by inputting gene symbol. Tab 2-CircPlotter) Query ERCC normalized BSJ data from the samples by inputting circBase annotation. Tab 3-Summary data) RNA Star 2-pass mapping log output file information, including input reads and those mapped to the genome assemblies. Tab 4-Raw counts) CircRNA counts are generated by CLEAR (CircExplorer3) mapping and include only BSJ reads. All other gene counts are outputs from RNA STAR GeneCount (per gene read counts). Tab 5-ERCC norm counts) BSJ or gene counts normalized using ERCC spike-in reads. Tab 6-Log2FC) $Log_2FC$ (Infected/Uninfected) for ERCC normalized counts.

**Supporting Dataset 6. Transcriptomic data from interferon stimulation experiments**
Transcriptomic data from bulk RNA-Seq data for MRC-5, LEC, or Akata- treated with IFN-$\beta$ and -$\gamma$ for 48 hours. Data was mapped to a concatenated genome assembly containing hg38 (gencode.v36) and ERCC spike-ins. Tab 1-GenePlotter) Query ERCC normalized gene expression data from the samples by inputting gene symbol. Tab 2-CircPlotter) Query ERCC normalized BSJ data from the samples by inputting circBase annotation. Tab 3-Summary data) RNA Star 2-pass mapping log output file information, including input reads and those mapped to the genome assemblies. Tab 4-Raw counts) CircRNA counts are generated by CLEAR (CircExplorer3) mapping and include only BSJ reads. All other gene counts are outputs from RNA STAR GeneCount (per gene read counts). Tab 5-ERCC norm counts) BSJ or gene counts normalized using ERCC spike-in reads. Tab 6-Log2FC) $Log_2FC$ (Infected/Uninfected) for ERCC normalized counts.

### SUPPLEMENTARY TABLES

#### Supplementary Table 1. Differentially expressed circRNAs

Summary table of all circRNAs which met our significance threshold to be called as differentially expressed. CircRNAs are annotated relative to their colinear gene and back splice junction and cross-referenced via circBase (human) or circAtlas (mouse).

#### Supplementary Table 2. Oligos used in this study

Sequences of oligos used for qPCR or other applications in this study.

| Target | Target Species | Forward Primer | Reverse Primer | Notes |
|---|---|---|---|---|
| **US1 (ICP22)** | HSV-1 | TTTGGGGAGTTTGACTGGAC | CAGACACTTGCGGTCTTCTG | |
| **UL23 (tk)** | HSV-1 | ACCCGCTTAACAGCGTCAACA | CCAAAGAGGTGCGGGAGTTT | Used to detect HSV-1 genome copies |
| **UL44 (gC)** | HSV-1 | GTGACGTTTGCCTGGTTCCTGG | GCACGACTCCTGGGCCGTAACG | |
| **circMAN1A2** | Human | TAGGACATATGGGTGGGGAC | TGCTTCTTCCAAGGCCTTCT | Divergent Primers (hsa_circ_0000116) |
| **circEPSTI1** | Human | AAGCTGAAGAAGCTGAACTC | GTGTATGCACTTGTGTATTGC | Divergent Primers (hsa_circ_0000479) |
| **circRELL1** | Human | ATGTCTGTTAGTGGGGCTGA | TATCTGCTACCATCGCCTTT | Divergent Primers (hsa_circ_0001400) |
| **circTNPO3** | Human | GCTAATCGGCGCACAGAAAT | GGTCTGAGATCTCCCATGCA | Divergent Primers (hsa_circ_0001741) |
| **circECE1** | Human | ACCTCTGGGAACACAACCAA | GCCGTTGGGGTATGCGTC | Divergent Primers (hsa_circ_0002402) |
| **circIPO7** | Human | GGGCCAGATGAAGAAGGTAGT | GAGCCTGCATTACAGGTCTGAT | Divergent Primers (hsa_circ_0005092) |
| ***RELL1*** | Human | GCAGTGGCACAGAGTAGCAG | CAGTGCAGCCTTACCAGTTG | Primers span exon-exon junction |
| ***EPSTI1*** | Human | GACAGAAGTGCCTGTCAAAGTG | GCCGTTTCAGTTCCAGTAATTC | Primers span exon-exon junction |
| ***ISG15*** | Human | GAGAGGCAGCGAACTCATCT | CTTCAGCTCTGACACCGACA | |
| **18S rRNA** | Human | GTAACCCGTTGAACCCCATT | CCATCCAATCGGTAGTAGCG | |
| ***GAPDH*** | Human | AGCCTCAAGATCATCAGCAATG | ATGGACTGTGGTCATGAGTCCTT | Primers span exon-exon junction |
| ***SHFL* (Shiftless)** | Human | GGCTCTGATGAGGAAATTCGGC | TCCTGGGCATCCAGATCGTTAC | |
| **U6 snRNA** | Human | GGAATCTAGAACATATACTAAAATTGGAAC | GGAACTCGAGTTTGCGTGTCATCCTTGCGC | |
| **5S rRNA** | Human | GGCCATACCACCCTGAACGC | CAGCACCCGGTATTCCCAGG | |
| **7SL** | Human | CAAAACTCCCGTGCTGATCA | GGCTGGAGTGCAGTGGCTAT | |
| ***MALAT1*** | Human | GTCATAACCAGCCTGGCAGT | GCTTATTCCCCAATGGAGGT | |
| ***ECE1* mRNA** | Human | CGGAGCACGCGAGCTATGA | CCGCCAGAAGTACCACCAAC | Primers span exon-exon junction |
| ***IPO7* mRNA** | Human | TCAATTTTGGAGGCCCAGCA | AGGTTGCATGATCAGGTCCC | Primers span exon-exon junction |
| ***MAN1A2*** | Human | CACATGATGATGTACAGCAGAGC | ATCGAACAGCAGGATTACCTGA | Primers span exon-exon junction |
| ***TNPO3*** | Human | ACATTGCAGCTCGTGTACCA | AGCATGACTCCACATCCTGC | Primers span exon-exon junction |
| ***GAPDH*** | Human | CAGAACATCATCCCTGCCTCTACT | GCCGAGCTTCCCGTTCA | Used to detect human genome copies |
| ***RPS13*** | Human | TCGGCTTTACCCTATCGACGCAG | ACGTACTTGTGCAACACCATGTGA | |
| **CpG DNA** | N/A | T*C*G*T*C*G*T*T*T*T*G*T*C*G*T*T*T*T*G*T*C*G*T*T (RNase Free HPLC Purification, * Phosphorothioate Bond) | | Used for CpG immunostimulation |

**Supplementary Table 3. RNA-Seq data analyzed in this study**

Overview of bulk RNA-Seq data analyzed in this study. If data from any of the sample sets was previously published we have included PubMed reference numbers.

| Dataset | Model | Experiment information | Replicates | Sequencing type | SRA or GEO Accession |
|---|---|---|---|---|---|
| **HSV-1 infection** | **Lytic infection** | MRC-5 cells infected with HSV-1 strain KOS (MOI 10 PFU/cell) for 12 or 24 hours. **PMID: 36724259** | 2-4 | Illumina 150 PE | SRR19779319, SRR19779318, SRR19787559 |
| | **Latent infection** | Female 8-week-old BALB/cAnTAC mice infected with HSV-1 strain 17 (10^5 PFU) via the ocular route. Latently infected trigeminal ganglion (TG) were harvested 4 weeks after primary infection. | 4 | Illumina 150 PE | SRR19792335, SRR19792334 |
| | **Lytic reactivation** | Female 8-week-old BALB/cAnTAC mice infected with HSV-1 strain 17 (10^5 PFU) via the ocular route. Trigeminal ganglia were explanted (TG explant) ~4 weeks after primary infection in the presence of vehicle or 2 uM JQ1 for 12 hours. | 2-3 | Illumina 150 PE | SRR25824398, SRR25824397, SRR25824394, SRR25824396 |
| **HCMV infection** | **Lytic infection** | MRC-5 cells infected with HCMV strain TB40/E (MOI 3 PFU/cell) for 24 or 72 hours. **PMID: 28874566** | 2 | Illumina 100 PE | SRR5629593, SRR5629594, SRR5629591, SRR5629592, SRR5629589, SRR5629590, SRR5629587, SRR5629588, SRR5629577, SRR5629578, SRR5629575, SRR5629576, SRR5629573, SRR5629574, SRR5629571, SRR5629572 |
| **KSHV infection** | **Lytic infection** | LEC infected with KSHV strain BAC16 (MOI 1 PFU/cell) for 24 or 72 hours. **PMID: 36724259** | 2 | Illumina 150 PE | SRR20020769, SRR25816558, SRR20020770 |
| | **Lytic reactivation** | iSLK-BAC16 induced with 1 ug/mL Doxycycline 1 mM Sodium Butyrate for 24 or 72 hours. **PMID: 36724259** | 4 | Illumina 150 PE | SRR25816557, SRR20020761, SRR25816556, SRR20020757, SRR20020758 |
| **MHV68 infection** | **Lytic infection** | NIH 3T3 cells infected with MHV68 strain H2B-YFP (MOI 5 PFU/cell) for 6 or 18 hours. | 2 | Illumina 150 PE | SRR19792326, SRR25823339, SRR19792325 |
| | **Lytic reactivation** | A20 HE-RIT cells induced with 5 ug/ml Dox and 20 ng/ml TPA for 6 or 24 hours. | 2 | Illumina 150 PE | SRR19792324, SRR25823338, SRR19792321 |
| **Interferon stimulation** | **Uninfected** | MRC-5, LEC, or Akata- cells treated with IFN-$\beta$ or -$\gamma$ or- for 48 hours. | 3 | Illumina 150 PE | SRR25905055, SRR25905049, SRR25905048, SRR25905050, SRR25905054, SRR25905051, SRR25905053, SRR25905052 |
| **EBV infection** | **Latent infection** | Akata+ (EBV positive) or Akata- (EBV negative) cells. 074301 Arraystar Human CircRNA microarray V2. **PMID: 36724259** | 3 | Microarray | GSE206824 |

### SUPPLEMENTARY FIGURES

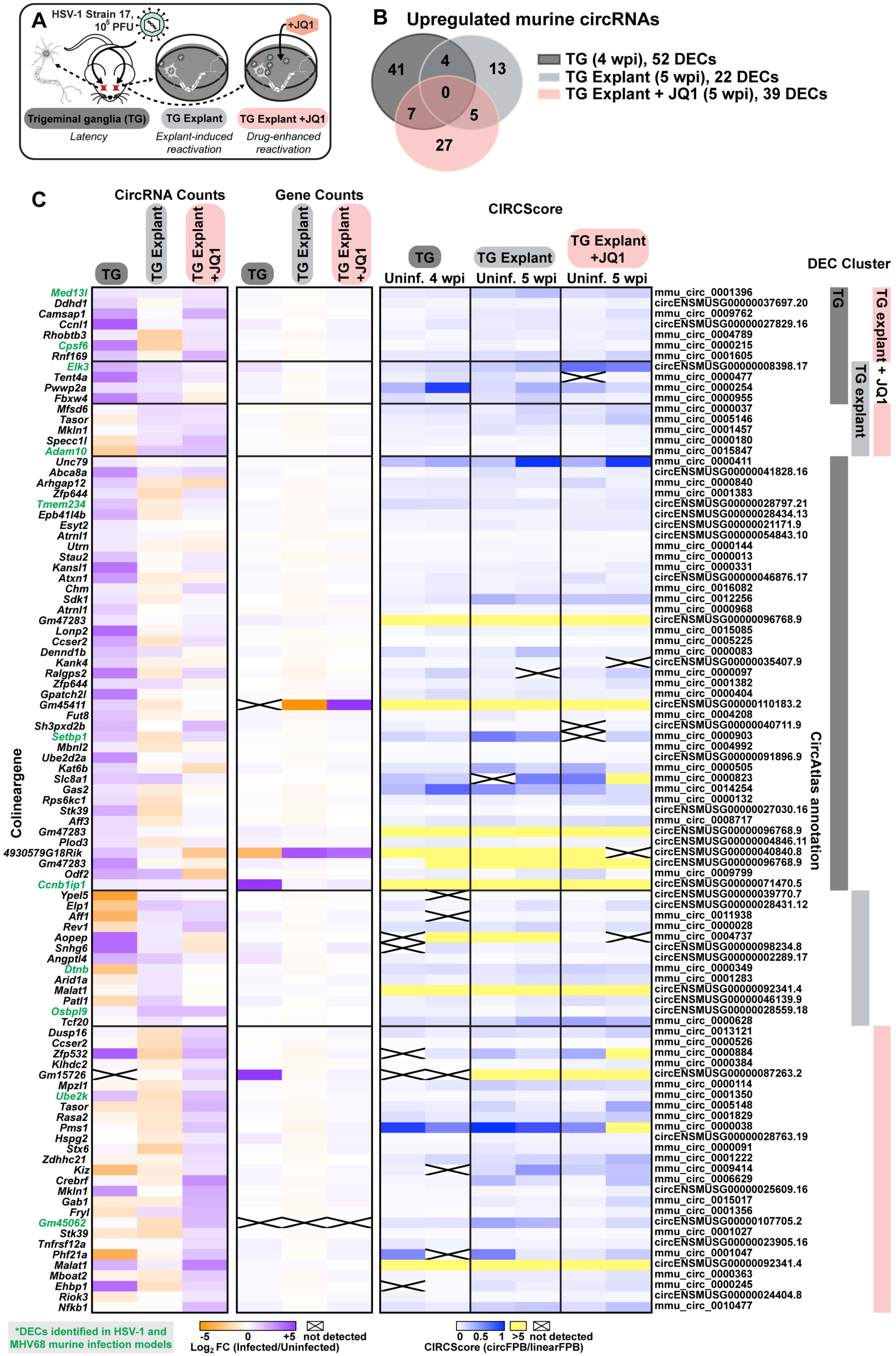

**Supplementary Figure 1-1. Murine circRNAs upregulated in HSV-1 infection models.**
A) Infographic for murine HSV-1 infection models used in this study. B) Overlap of upregulated murine circRNA detected by bulk RNA-Seq from HSV-1 latency (TG, n=4), explant-induced reactivation (TG explant, n=3), and drug-enhanced reactivation (TG explant + JQ1, n=2). Upregulated circRNAs had a raw BSJ count across the sample set >10, $Log_2FC>0.5$, and RankProduct p-value <0.05. C) Heatmaps for upregulated murine circRNAs, plotted as circRNA counts ($Log_2FC$ Infected/Uninfected ERCC normalized BSJ counts), Gene counts ($Log_2FC$ Infected/Uninfected ERCC normalized gene counts), or CIRCScore (circFPB/linearFPB). Heatmap values are the average of biological replicates. $Log_2FC$ is relative a paired uninfected control. Differentially expressed circRNAs (DECs) from murine HSV-1 and MHV68 infection models have their colinear gene names colored green (11 overlapping DECs).

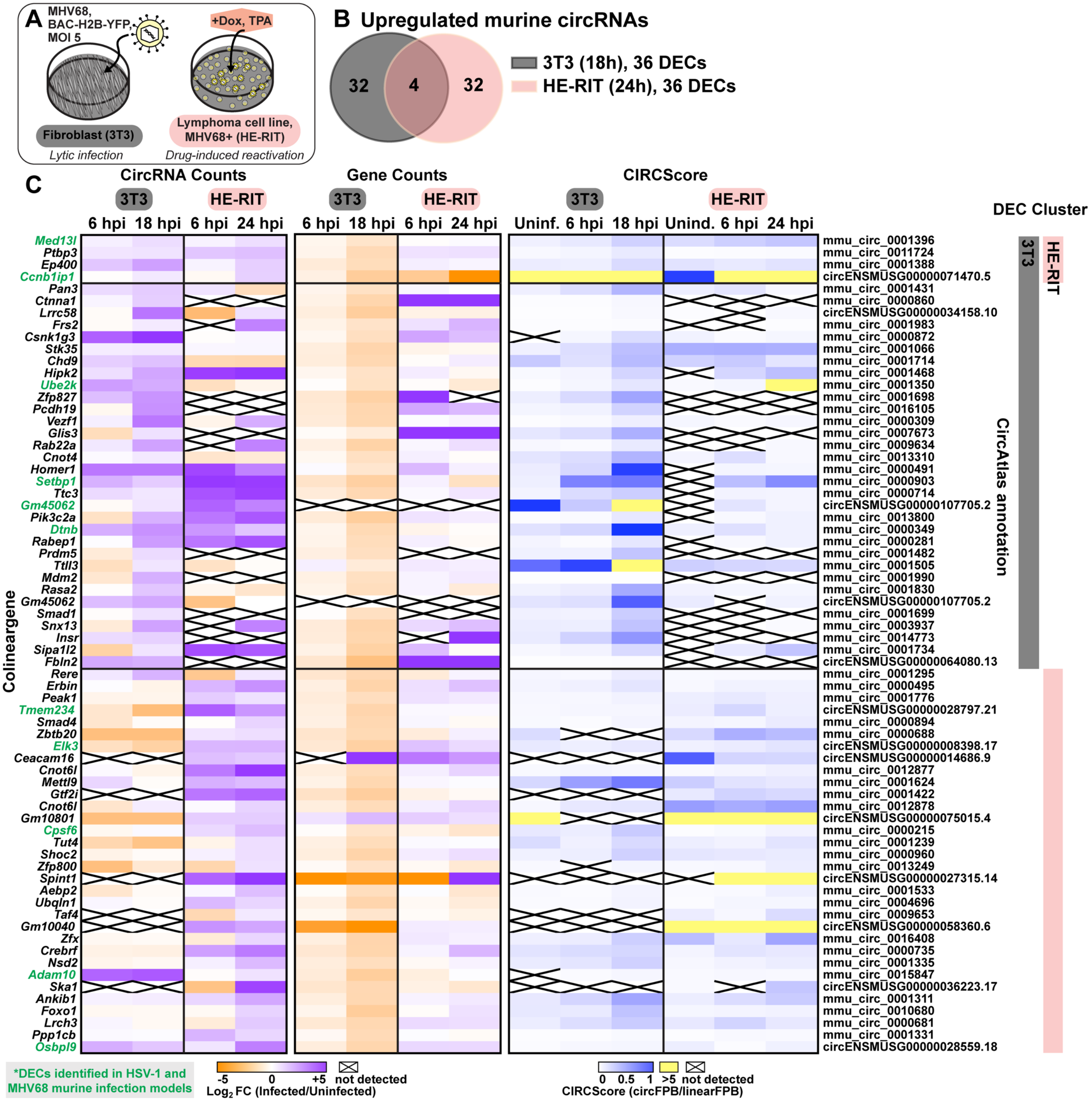


**Supplementary Figure 1-2. Murine circRNAs upregulated in MHV68 infection models.**
A) Infographic for murine MHV68 infection models used in this study. B) Overlap of upregulated murine circRNA detected by bulk RNA-Seq from MHV68 *de novo* infection (NIH 3T3, n=2) and drug-induced reactivation (HE-RIT, n=2). Upregulated circRNAs had a raw BSJ count across the sample set >10, $Log_2FC>0.5$, and p-value <0.05. C) Heatmaps for upregulated murine circRNAs, plotted as CircRNA counts (3T3: $Log_2FC$ Infected/Uninfected ERCC normalized BSJ counts, HE-RIT: $Log_2FC$ Drug treated/Uninduced ERCC normalized BSJ counts), Gene counts (3T3: $Log_2FC$ Infected/Uninfected ERCC normalized BSJ counts, HE-RIT: $Log_2FC$ Drug treated/Uninduced ERCC normalized BSJ counts), or CIRCScore (circFPB/linearFPB). Heatmap values are the average of biological replicates. $Log_2FC$ is relative a paired uninfected or uninduced control. Differentially expressed circRNAs (DECs) from murine HSV-1 and MHV68 infection models have their colinear gene names colored green (11 overlapping DECs).

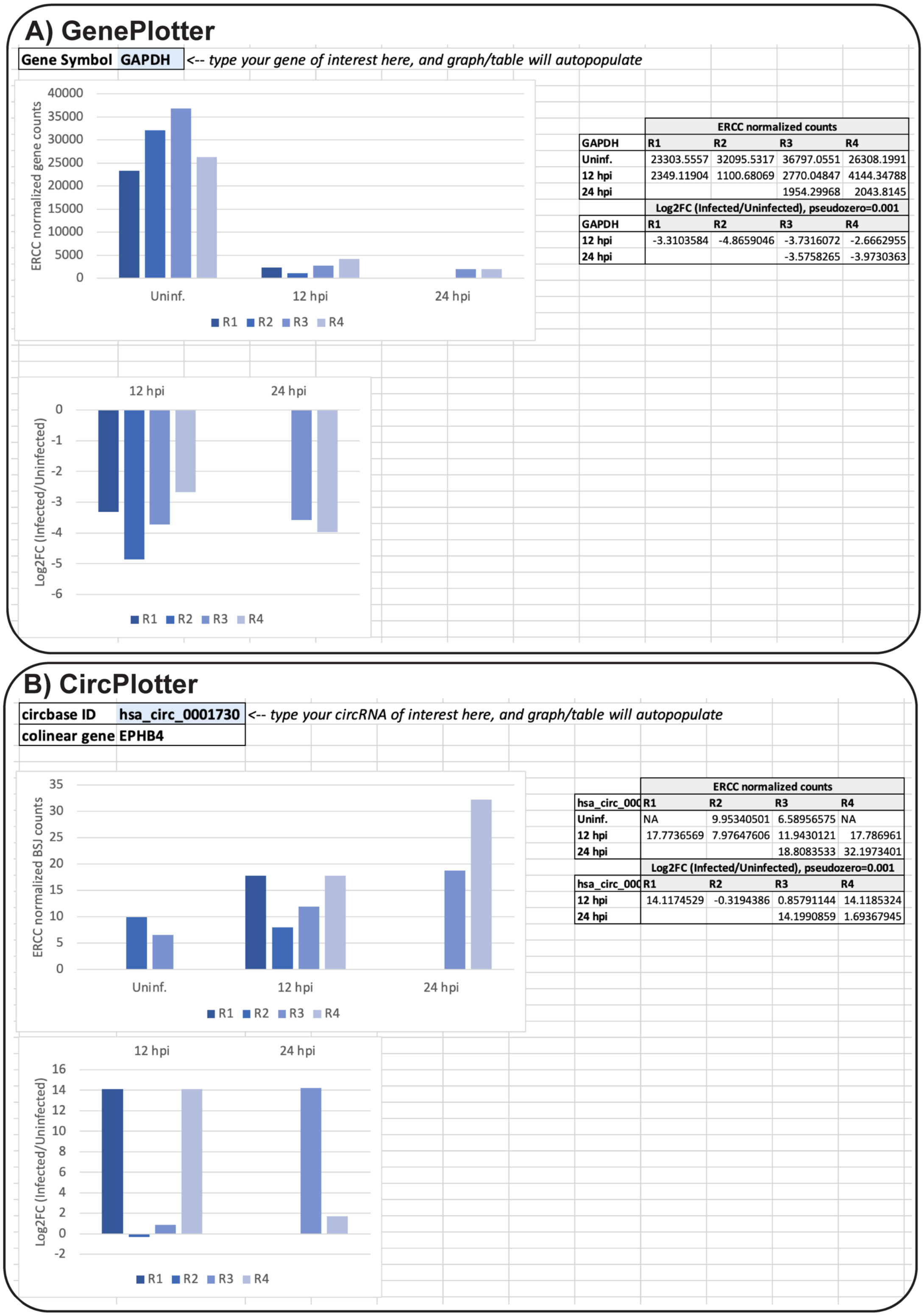


**Supplementary Figure 1-3. Screenshot of interactive resource table for transcriptomic data.** Supporting datasets were generated for all bulk RNA-Seq data analyzed in this study. Datasets include raw and normalized gene and BSJ counts. Users can query any gene or circRNA of interest using the GenePlotter of CircPlotter tabs.

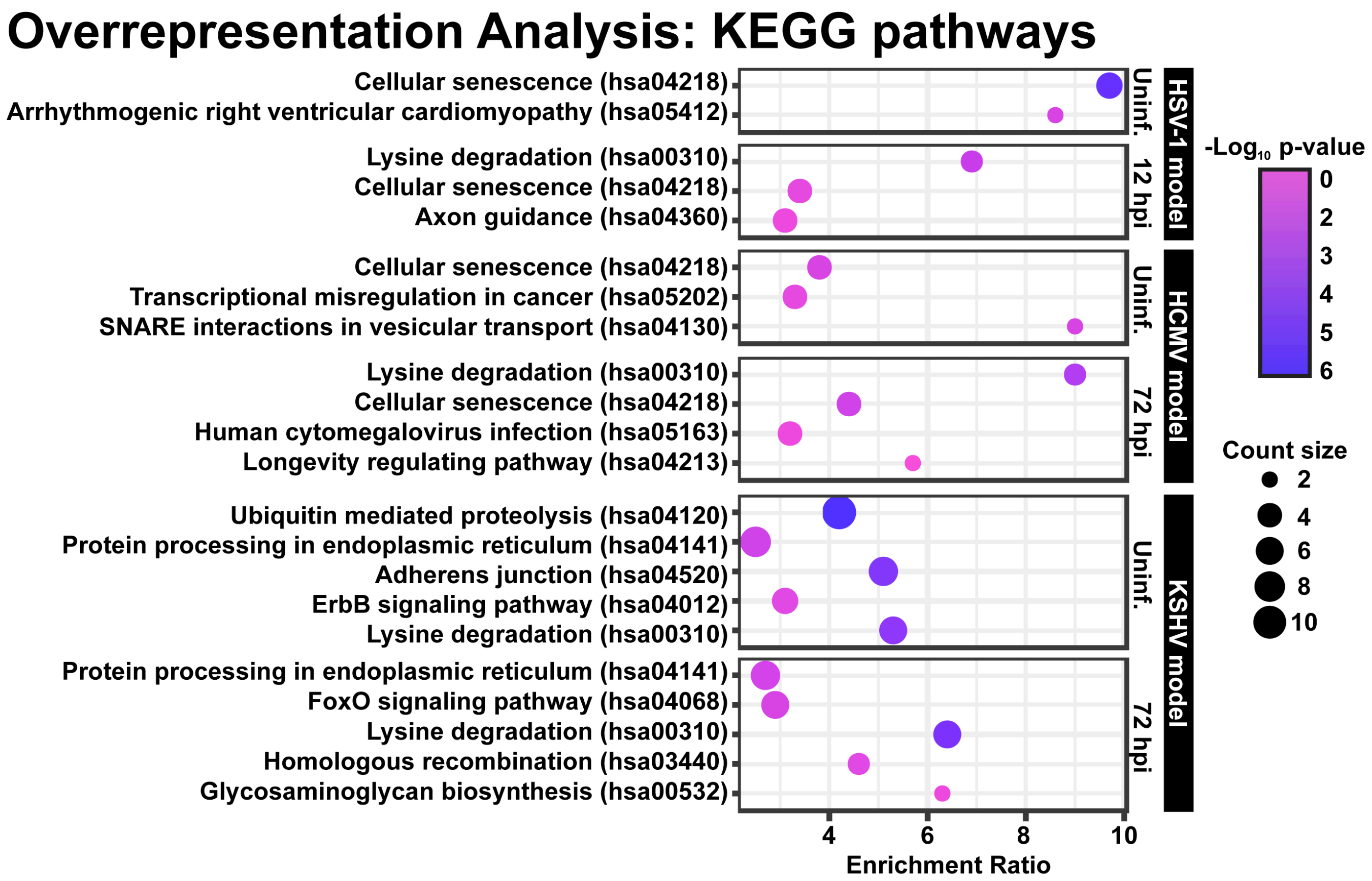


**Supplementary Figure 1-4. Overrepresentation analysis (ORA) of colinear genes containing circRNAs expressed in human herpesvirus infection models.** ORA was performed using a list of genes colinear to circRNAs detected in a given model (>10 raw BSJ read counts). Data is represented as a pathway enrichment bubble plot, with bubble size indicating the number of genes from a given pathway, y-axis is enrichment ratio (relative to expected from a random gene sampling), and bubble color is -Log$_{10}$ p-value. Only pathways with a p-value of <0.05 were included.

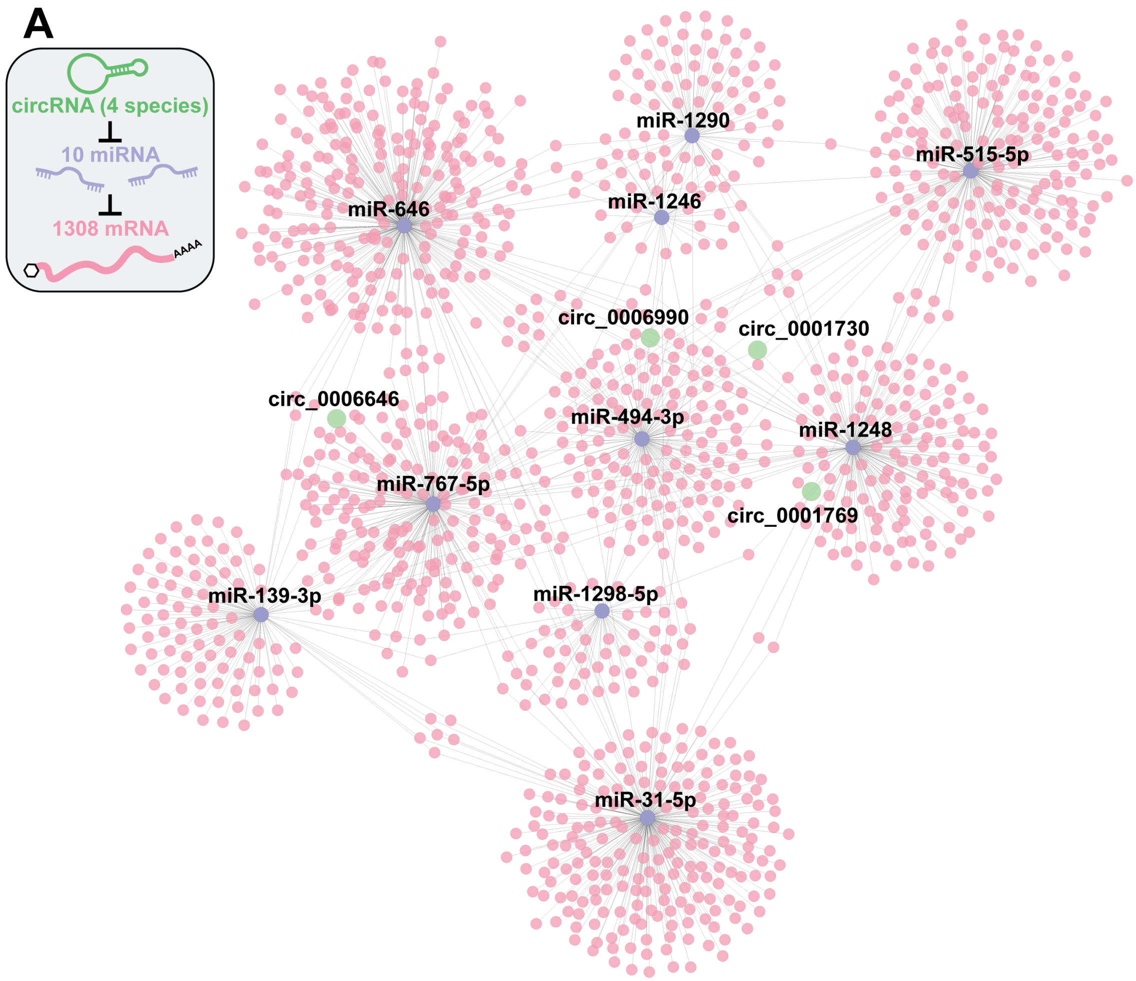


**B miEAA-miRNA Enrichment Annotation**

**MHC class I protein complex assembly (GO0002397), p-value=2.06e-5**
hsa-miR-1248; hsa-miR-515-5p; hsa-miR-1246; hsa-miR-1290; hsa-miR-1298-5p; hsa-miR-646; hsa-miR-31-5p

**MHC class Ib protein complex assembly (GO0002398), p-value=2.06e-5**
hsa-miR-1248; hsa-miR-515-5p; hsa-miR-1246; hsa-miR-1290; hsa-miR-1298-5p; hsa-miR-646; hsa-miR-31-5p

**TAP2 binding (GO0046979), p-value=3.33e-5**
hsa-miR-1248; hsa-miR-515-5p; hsa-miR-1246; hsa-miR-1290; hsa-miR-1298-5p; hsa-miR-646; hsa-miR-31-5p

**TAP complex binding (GO0062061), p-value=3.35e-5**
hsa-miR-1248; hsa-miR-515-5p; hsa-miR-767-5p; hsa-miR-1246; hsa-miR-1290; hsa-miR-1298-5p; hsa-miR-139-3p; hsa-miR-646; hsa-miR-31-5p

**Tapasin-ERp57 complex (GO0061779), p-value=2.57e-5**
hsa-miR-1248; hsa-miR-515-5p; hsa-miR-1246; hsa-miR-1290; hsa-miR-1298-5p; hsa-miR-494-3p; hsa-miR-646; hsa-miR-31-5p

**Peptide antigen stabilization (GO0050823), p-value=2.06e-5**
hsa-miR-1248; hsa-miR-515-5p; hsa-miR-1246; hsa-miR-1290; hsa-miR-1298-5p; hsa-miR-646; hsa-miR-31-5p

**Supplementary Figure 1-5. In silico network predictions for circRNA-miRNA-mRNA networks.** A) Four commonly-regulated human circRNAs (hsa_circ_0001730, hsa_circ_0006990, hsa_circ_0006646, hsa_circ_0001769) were examined for miRNA-binding sites using CircInteractome (1). Experimentally supported miRNA-mRNA interactions were then added to the network based on DIANA-TarBase v8.0 (2). Cytoscape 3.10 was used for visualizing the network. B) Overrepresentation analysis of predicted target miRNAs was performed with miEAA2.0 (4). Enriched Gene Ontology terms (http://geneontology.org), p-values, and miRNAs that were predicted to be regulated by circRNAs are shown.

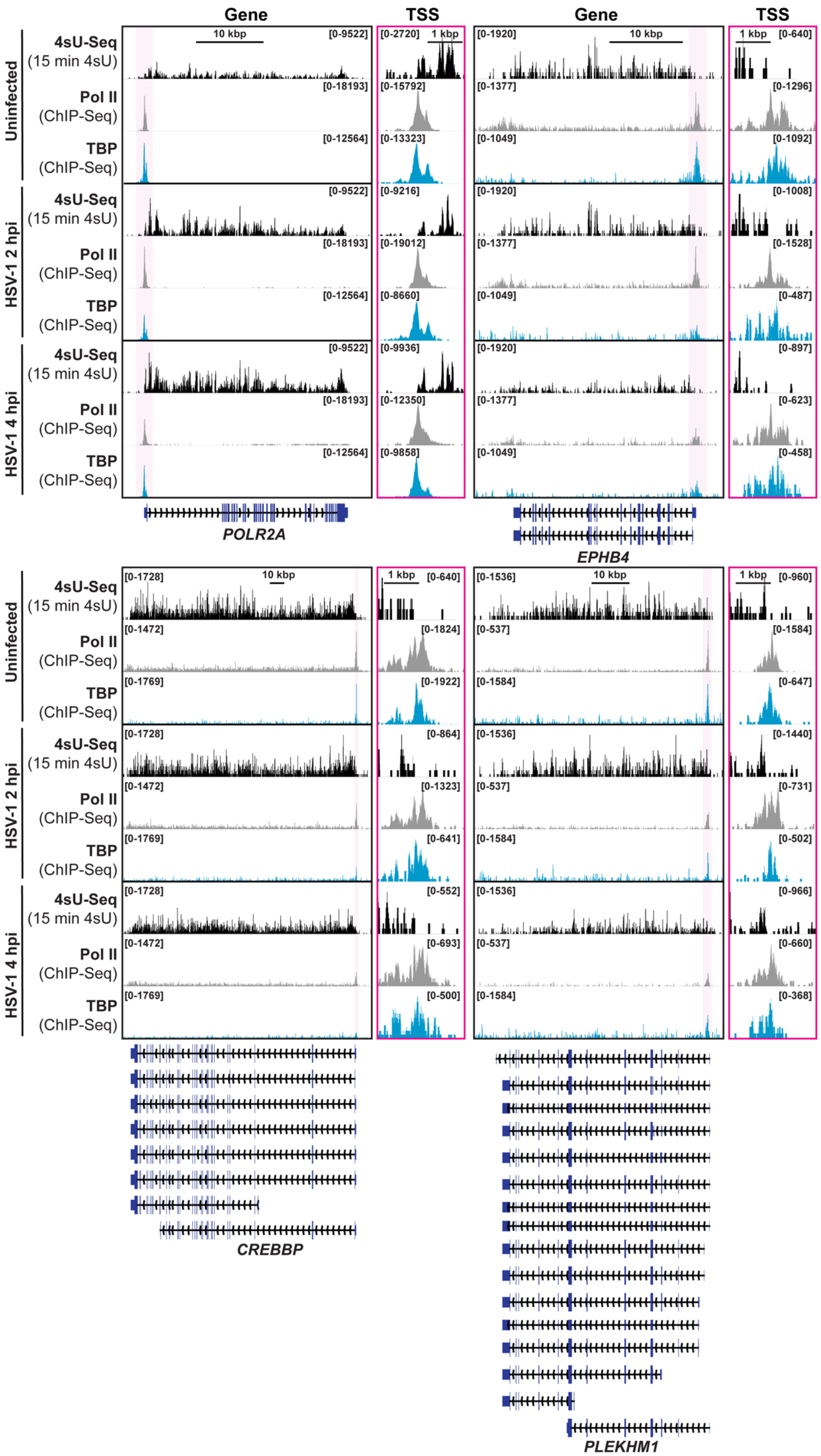
Gene
TSS
Gene
TSS
10 kbp
1 kbp
Uninfected
HSV-1 2 hpi
HSV-1 4 hpi
4sU-Seq
(15 min 4sU)
Pol II
(ChIP-Seq)
TBP
(ChIP-Seq)
POLR2A
EPHB4
CREBBP
PLEKHM1

**Supplementary Figure 3-1. Transcriptional activity of genes containing circRNAs upregulated after HSV-1 infection.** Nascent transcription and binding of general transcription factors for genes that host herpesvirus infection-induced circRNAs. Our previously published data from fibroblasts infected with HSV-1 (MRC-5 + KOS MOI 10) was reanalyzed (8). 4sU-Seq data is normalized to rRNA mapped reads. ChIP-Seq data is the average of biological duplicates and normalized to sequencing depth. Y-axes maximum and minimum values are listed within brackets. Traces spanning genes have the same y-axis maximums across infection conditions. Traces spanning 2.5 kbp transcription start sites (TSS) have autoscaled y-axis maximums.

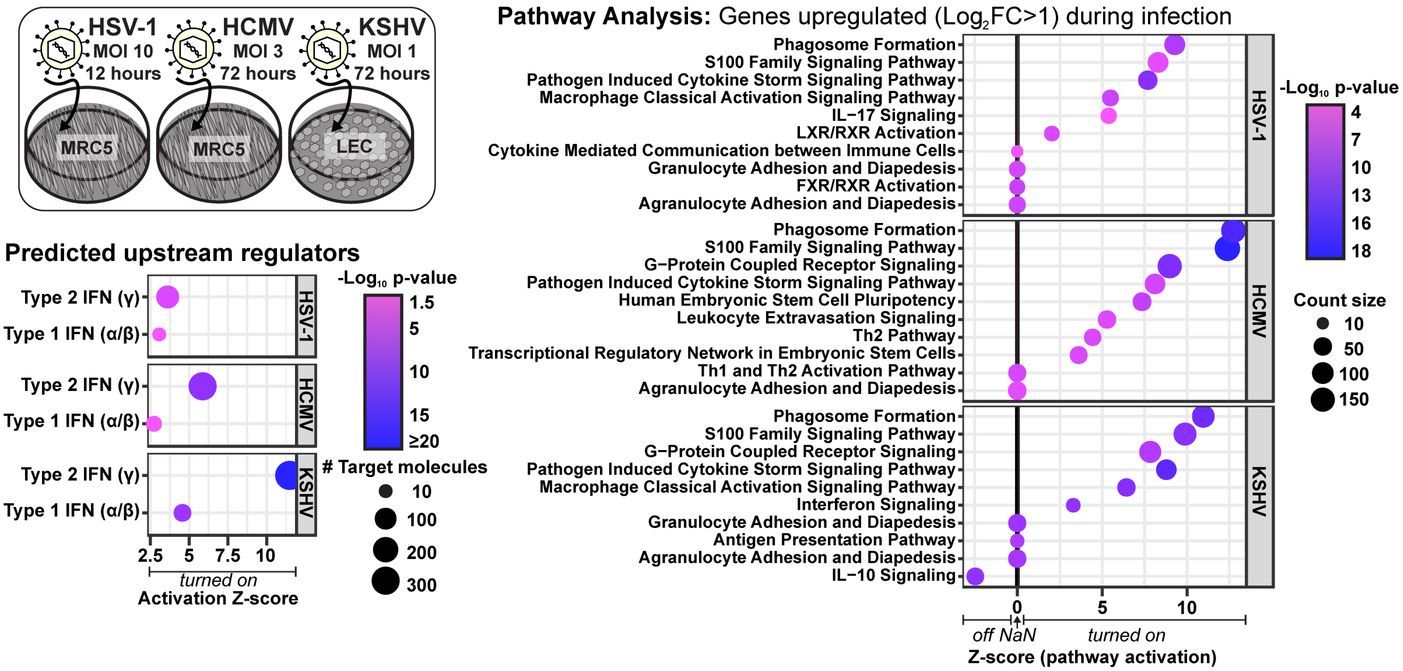


**Supplementary Figure 4-1. Ingenuity pathway analysis (IPA) for upregulated genes during herpesvirus lytic infection.** IPA (Qiagen) was performed on bulk RNA-Seq datasets for HSV-1 (n=4), HCMV (n=2), and KSHV (n=2). RNA-Seq data was normalized using ERCC spike-in controls. Only genes with an average $Log_2FC$ >1 (Infected/Uninfected) were used for IPA. The predicted upstream role of Type I and II interferon based on upregulated gene identities was presented in a bubble plot. Z-score indicates activation status, -$log_{10}$p-value indicates enrichment significance, and count size is the number of target molecules downstream of IFN that were upregulated. The top 10 most significantly (by p-value) altered pathways were plotted. Z-score indicates predicted pathway activation, -$log_{10}$p-value indicates enrichment significance, and count size is the number of upregulated genes in the pathway.

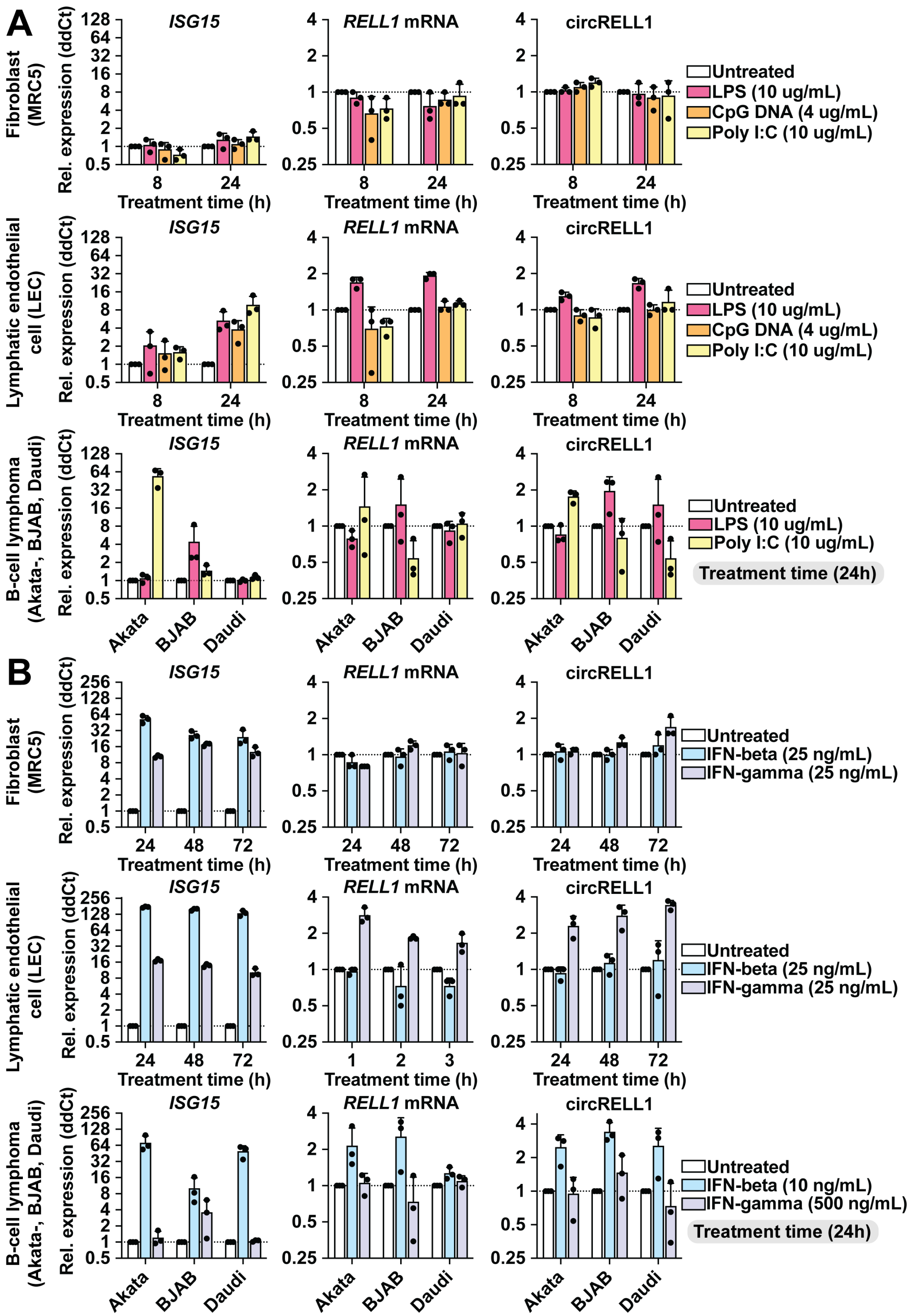


**Supplementary Figure 4-2. CircRELL1 expression changes in response to immune stimuli.** Fibroblasts (MRC-5), lymphatic endothelial cells (LEC), or B-cell lymphoma cells (Akata-, BJAB, Daudi) were treated with immune stimulants including A) lipopolysaccharide (LPS), poly I:C, CpG DNA, or B) IFN-$\beta$ and -$\gamma$. RNA was collected and assessed after reverse transcription using qPCR. Data is plotted as relative expression (ddCt) using 18S rRNA as the reference gene, and relative to a paired untreated sample. Data points are biological replicates, column bars are the average, and error bars are standard deviation.

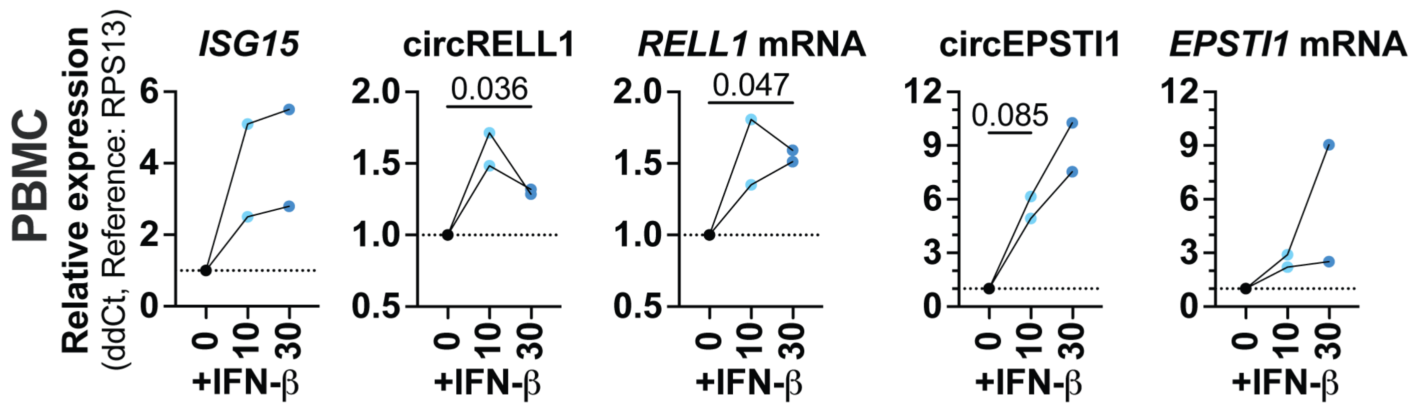


**Supplementary Figure 4-3. Interferon stimulated circRNAs in PBMCs.**
Human peripheral blood peripheral blood mononuclear cells (PBMCs) were isolated from two donors and treated with IFN-β (0, 10, or 30 μg/ml) for 24 hours. RNA was collected and assessed after reverse transcription using qPCR. Data is plotted as relative expression (ddCt) using *RPS13* mRNA as the reference gene, and relative to a paired untreated sample. Data points from same donors are connected by lines. Significances were calculated using paired t-tests and any p-values < 0.1 are shown.
